# Single cell transcriptomics reveals opioid usage evokes widespread suppression of antiviral gene program

**DOI:** 10.1101/630574

**Authors:** Tanya T. Karagiannis, John P. Cleary, Busra Gok, Nicholas G. Martin, Elliot C. Nelson, Christine S. Cheng

## Abstract

Chronic opioid usage not only causes addiction behavior through the central nervous system (CNS), but it also modulates the peripheral immune system. However, whether opioid usage positively or negatively impacts the immune system is still controversial. In order to understand the immune modulatory effect of opioids in a systematic and unbiased way, we performed single cell RNA sequencing (scRNA-seq) of peripheral blood mononuclear cells (PBMCs) from opioid-dependent individuals and non-dependent controls. We show that chronic opioid usage evokes widespread suppression of interferon-stimulated genes (ISGs) and antiviral gene program in naive monocytes and upon ex vivo stimulation with the pathogen component lipopolysaccharide (LPS) in multiple innate and adaptive immune cell types. Furthermore, scRNA-seq revealed the same phenomenon with in vitro morphine treatment; after just a short exposure to morphine stimulation, we observed the same suppression of antiviral genes in multiple immune cell types. These findings indicate that both acute and chronic opioid exposure may be harmful to our immune system by suppressing the antiviral gene program, our body’s defense response to potential infection. Our results suggest that further characterization of the immune modulatory effects of opioid use is critical to ensure the safety of clinical opioid usage.

---

The opioid epidemic is a major threat to global public health that affects millions of people and their families. Part of the problem is caused by the rapid increase in the number of opioid prescriptions written by medical practices starting from the late 1990s. From 1999 to 2017, overdoses related to prescription opioids have dramatically increased in the United States with overdose deaths found to be five times higher in 2017 compared to 1999^1^. In addition, opioids affect not only the CNS but also the peripheral immune system through the expression of a variety of opioid receptors on different immune cell types^2^. However, the effect of opioids on the peripheral immune system is complicated and involves various mechanisms. Studies have shown inconsistent results, where some suggest opioid usage is immunosuppressive while, in contrast, others suggest opioids are immunoactivating^2–4^. Most of these studies focus on a particular immune cell subpopulation and a few candidate genes. Interestingly, epidemiological studies suggest that opioid usage is associated with increased susceptibility to opportunistic infections such as tuberculosis, HIV and pneumonia^5–7^.

Given the extensive use of prescription opioids and the global opioid epidemic, it is important to understand how opioid usage modulates different cell types in the immune system. Primary cells such as peripheral blood immune cells are usually comprised of heterogeneous cell populations. It is therefore highly time-consuming and labor-intensive to separate and study the individual cell types and generally not feasible given the limited input material from patient samples. The recent development of droplet-based single cell RNA-sequencing (scRNA-seq) approaches allows profiling of gene expression of thousands of single cells from a limited quantity of heterogeneous patient samples^8, 9^. Here, we utilize droplet-based scRNA-seq to systematically characterize cell type specific gene expression in the peripheral immune system of opioid-dependent individuals compared to controls.

Using droplet-based scRNA-seq, we profiled gene expression in 57,271 single cells from the peripheral blood mononuclear cells (PBMCs) of 7 opioid-dependent individuals and 7 age/sex-matched non-dependent controls (averaging 3,980 single cells per individual) (Figure 1A). To examine opioid usage-specific changes in gene expression in response to pathogenic stimuli, we stimulated PBMCs from 3 of the 7 opioid-dependent individuals and 3 of the controls with lipopolysaccharide (LPS, a component of gram-negative bacteria) for 3 hours and profiled 22,326 single cells. We sequenced these single cells to an average depth of 21,801 reads per cell and detected on average 805 genes and 2,810 transcripts per cell. To identify each of the immune cell subpopulations, we applied dimensionality reduction methods, including principal component analysis (PCA) and t-stochastic neighbor embedding (t-SNE), and unsupervised graph-based clustering^10, 11^. We identified 12 immune cell types/states using expression of canonical gene markers (Figure 1B, Figure S1-S3). Of the naive state immune populations, we identified CD4+ T cells, naïve CD8+ T cells, memory CD8+ T cells, NK cells, B cells, and monocytes. Of the LPS-stimulated immune populations, we identified CD4+ T cells, CD8+ T cells, activated T cells, B cells, NK cells, and monocytes. (Figure 1B, Figure S1-S3). We observed a slight shift in global gene expression states between opioid-dependent and control samples in most of the naïve cell types, while we observed larger differences in gene expression states in LPS-stimulated cell types (Figure 1B).

**Figure 1.**
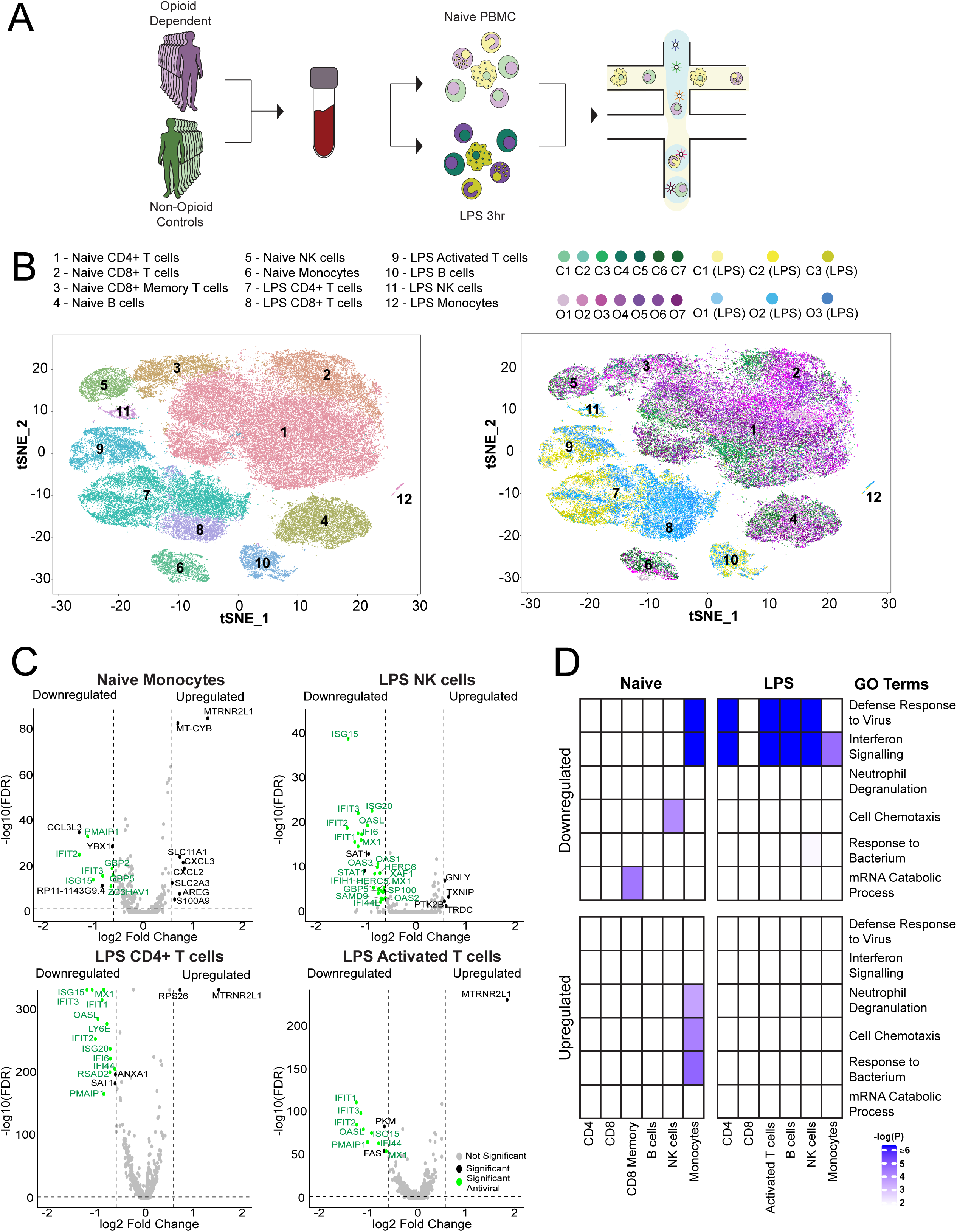
Single cell transcriptomics of PBMCs from opioid-dependent individuals revealed a widespread suppression of interferon-stimulated and antiviral genes upon LPS stimulation. **a,** Experimental workflow schematic. Peripheral blood from opioid-dependent individuals and control individuals were collected, PBMCs were isolated, and droplet based scRNA-seq was performed using Chromium Controller (10X Genomics). b, t-SNE plot of naive (51,336) and LPS (100ng/mL) stimulated (21,878) PBMCs were clustered (cells were filtered based on >300 and <2000 genes per cell, < 10000 UMIs per cell; see Methods) and identified into immune populations (top) and visualized by control individuals and opioid-dependent individuals in each state (bottom): naive state control samples 1-7 (C1-C7), naive state opioid-dependent samples 1-7 (O1-O7), LPS-stimulated control samples 1-3 (C1-C3 (LPS)), LPS-stimulated opioid-dependent samples 1-3 O1-O3 (LPS). c, Volcano plot showing fold change of gene expression (log2 scale) for downregulated and upregulated genes for opioid-dependent cells compared to non-dependent controls in naive monocytes, LPS stimulated NK cells, LPS stimulated CD4+ T cells, and LPS stimulated activated T cells (x-axis) with a significance of 0.05 (y-axis, −log10 scale). Significant genes shown in black, significant antiviral genes shown in green, and insignificant genes shown in grey. d, Pathway enrichment analysis of significant differential genes across all naive and LPS stimulated cell types evaluated by −log10(P-value) as indicated by blue-purple scale (x-axis:cell type/state, y-axis: pathways). White represents an analysis which did not provide enrichment results for the specific pathway.

From differential gene expression analysis for each immune subpopulation, we observed a down-regulation of interferon-stimulated genes and antiviral genes in opioid-dependent samples compared to control samples. The suppression of antiviral genes was observed only in monocytes in naive state and in all immune cell subpopulations under LPS stimulation except for CD8+ T cells and monocytes (Figure 1C, Figure S4-S5). This was further confirmed with gene set enrichment analysis of the resulting differential genes. We observed higher enrichment of defense response to virus and interferon signalling pathways in monocytes in naïve state and in most of the immune cell subpopulations in LPS-stimulated state (Figure 1D). When focusing on the three major innate immune response transcriptional modules activated by LPS stimulation^12^ (Figure 2A, Supplementary Table 1), we found widespread suppression of antiviral genes in opioid-dependent cells across LPS-stimulated immune subpopulations while peaked and sustained inflammatory genes were modestly affected by opioid usage (Figure 2B, Figure S6-S10). Given that antiviral genes are activated by an LPS induced type I interferon (IFNα and IFNβ) autocrine loop^13^, the potential mechanism for the suppression of antiviral genes in opioid users could be a higher expression of negative regulators of the interferon signaling pathway. Interestingly, we found the expression of two major interferon signaling inhibitors, suppressor of cytokine signaling *SOCS3*^14^ and ubiquitin-specific peptidase *USP15*^15^, to be upregulated in opioid-dependent cells in the naïve state: CD4+ T cells and monocytes for *SOCS3* and NK cells, CD8+ T cells and B cells for *USP15* (Figure S11). Taken together, our data suggest that chronic opioid usage results in widespread suppression of antiviral genes affecting all immune subpopulations including both innate and adaptive immune cell types.

**Figure 2.**
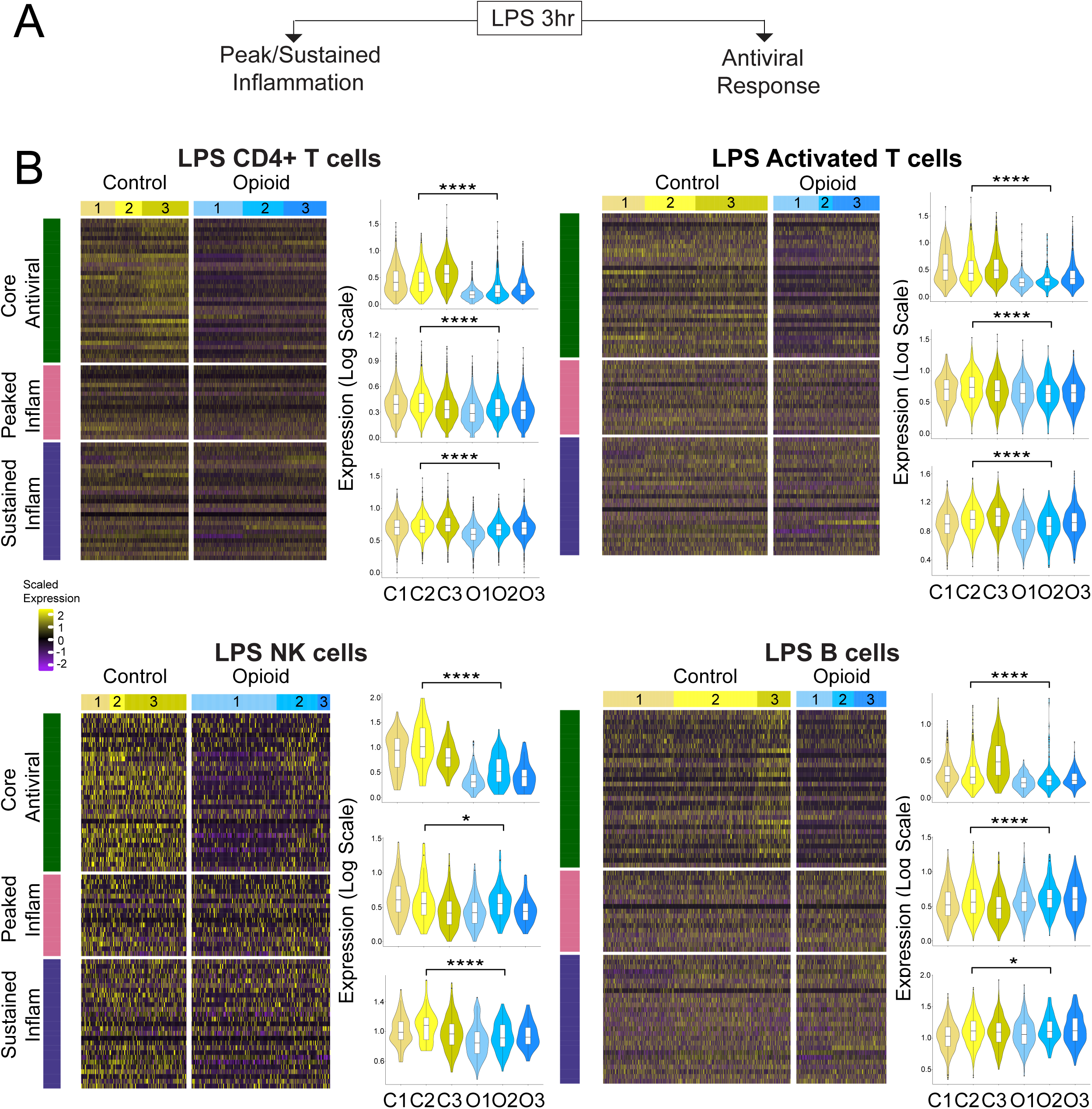
LPS-stimulated antiviral gene program showed consistent suppression in each of the immune cell populations from opioid dependent individuals. **a,** Evaluation of the three innate immune response gene programs stimulated by LPS: antiviral, peaked inflammatory, and sustained inflammatory. **b,** Heatmap of scaled expression of core antiviral and inflammatory response genes (left) and average gene set expression (log expression) (right) observed in control and opioid-dependent cells of LPS-stimulated samples: control samples 1-3 (C1-C3) and opioid-dependent samples 1-3 (O1-O3). scRNA-seq gene expression profiles were from PBMCs stimulated with LPS in CD4+ T cells, activated T cells, NK cells, and B cells. We found significant suppression of core antiviral gene expression from opioid-dependent individuals compared to control individuals (**** (p-value) < 0.0001). The three core innate immune response gene modules:^12^: core antiviral (Core Antiviral), peaked inflammation (Peaked Inflam), and sustained inflammation (Sustained Inflam) (Supplementary Table 1).

To examine the in vitro effect of opioids, we first treated primary human PBMCs from healthy individuals with a titration of morphine for 24 hours before stimulating with either a mock treatment (Untreated) or 100ng/mL LPS for three hours. We then performed RT-qPCR using primers against the major antiviral gene, *ISG15*, which was was the most prominent antiviral gene downregulated in opioid-dependent cells across cell types. We found that PBMCs pretreated with morphine for 24 hours exhibited a dose-dependent inhibition to the induction of *ISG15* after LPS stimulation (Figure 3A). Furthermore, this inhibition was observed as quickly as three hours after morphine pretreatment (Figure 3B). In order to characterize this phenomenon at a genome-wide scale and in every immune cell type in a cost-effective way, we performed scRNA-seq with an antibody-based cell hashing technique^16^ to multiplex samples in droplet-based scRNA-seq (Figure 3C, Figure S12; see Methods). We profiled 2,958 single PBMCs treated with morphine alone and then stimulated with LPS for 3 hours. We found a modest but consistent suppression of core antiviral genes in response to morphine exposure. This phenotype was most pronounced in CD4+ T cells, CD8+ T cells, and NK cells (Figure 3C; Figure S13-S17).

**Figure 3:**
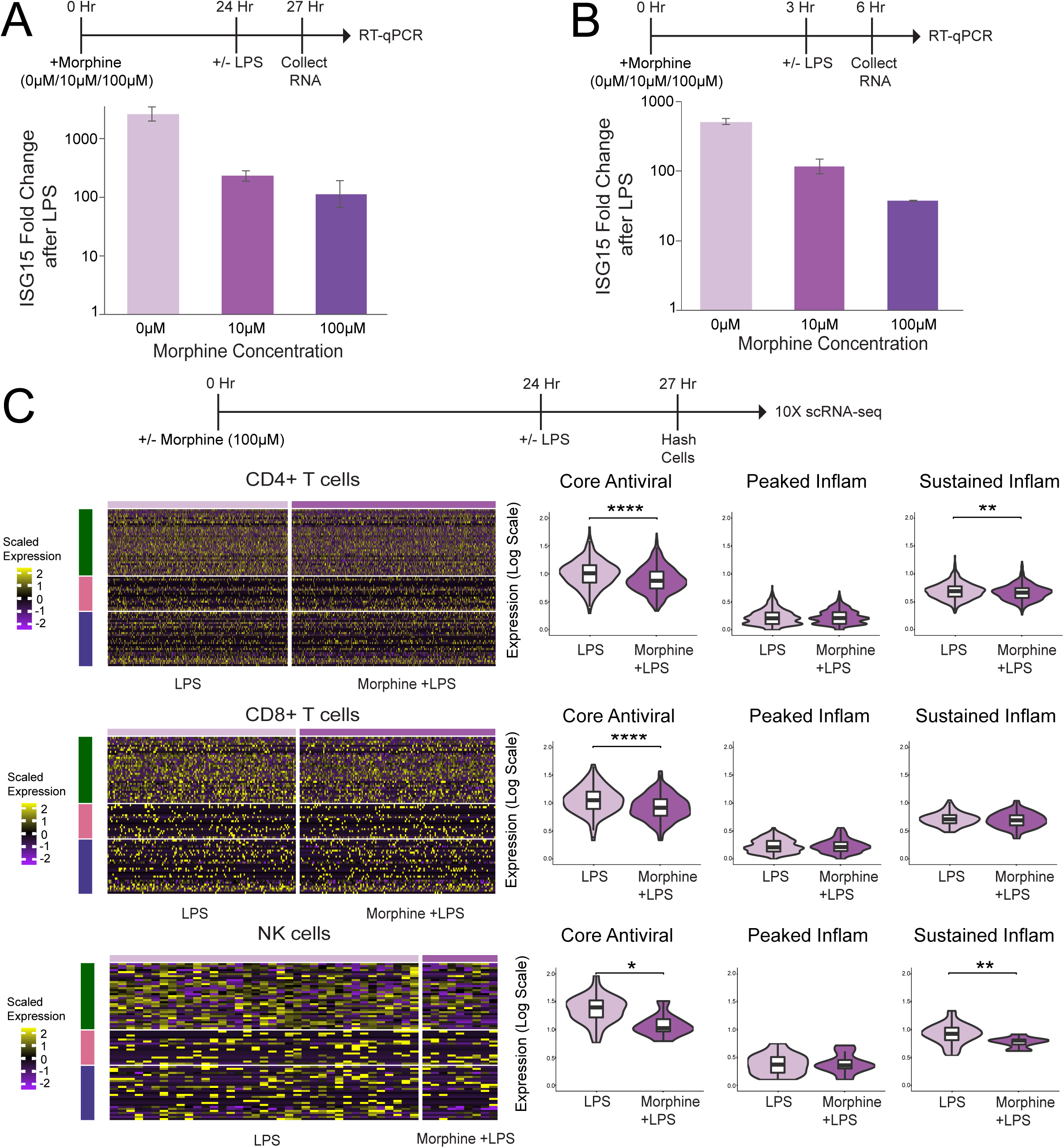
A short exposure to morphine treatment resulted in suppression of antiviral genes upon LPS stimulation. **a,** Evaluation of mRNA expression level of ISG15 upon dose response to morphine. PBMCs from a healthy individual were pre-treated with a titration of morphine (0μM, 10μM, 100μM) for 24 hours followed by LPS (100ng/mL) stimulation for 3 hours. Expression of interferon pathway gene ISG15 was evaluated by RT-qPCR. Values are displayed as fold increase (log10) to gene expression in LPS stimulated cells over unstimulated cells. **b,** Cell hashing scRNA-seq of healthy PBMCs pre-treated with morphine as described followed by LPS (100ng/mL) stimulation for 3 hours. Heatmaps of scaled single cell expression value of core antiviral genes (left) and violin plots of average core antiviral gene set expression (log scaled) (right) observed in LPS stimulated cells (LPS), morphine treated and LPS stimulated cells (Morphine + LPS) in CD4+ T cells, CD8+ T cells, and NK cells. Significance shown between LPS and Morphine + LPS (**** (p-value) < 0.0001, * (p-value) < 0.05).

Our results show that there is widespread suppression of interferon-stimulated genes and antiviral genes in multiple innate and adaptive peripheral immune subpopulations both ex vivo and in vitro upon LPS stimulation. Our findings suggest a potential adverse effect from opioid usage on the defense response towards viral infection in the immune system. This may explain in part the higher susceptibility to viral infection in opioid users observed in epidemiological studies^5–7, 17^. In addition, our in vitro findings also demonstrate that the observed suppression of antiviral pathway from our ex vivo experiments does not arise from needle sharing or presence of hepatitis C virus (HCV) infection in these injection opioid users. Given that most recreational opioid users inject drugs and many prescription opioid users are post-surgery or cancer patients under chemotherapy treatments, they are already more prone to infection; therefore suppression of antiviral genes with opioid usage brings clinical relevance and demonstrates the importance of carefully examining each individual case to avoid any possibility of comorbidity.

Opioid-induced immune modulation is mainly thought to occur through opioid receptors present on peripheral immune cell types^2, 18–20^. However, the presence of opioid receptors in peripheral immune cells is controversial. While several studies have shown that classical opioid receptors such as MOP are expressed on various peripheral blood immune cell types^21–26^, other studies evaluated the presence of opioid receptors in PBMCs in which they failed to detect mRNA transcripts for all opioid receptors except for the nonclassical receptor NOP^4, 27^. To clarify this, we looked at the expression of opioid receptors in population level RNA-seq data from PBMCs of healthy individuals from a previous study^28^. We found that the classical opioid receptor MOP is expressed in CD4+ T cells, CD8+ T cells, monocytes and NK cells, but not in B cells, while the other two classical opioid receptors DOP and KOP are very low in expression or undetectable (Supplementary Figure S18). The nonclassical receptor NOP is expressed in all immune cell types and is higher expressed in monocytes (Supplementary Figure S18). We anticipate the immune modulatory effect we observed from in vivo opioid usage and in vitro opioid treatment potentially occurs through both MOP and NOP receptors.

The stimulation of toll-like receptor 4 *TLR4* by LPS induces expression of hundreds of innate immune response genes previously categorized into three gene modules: antiviral, peaked inflammatory and sustained inflammatory genes^12^. Type I interferons function as autocrine and paracrine factors to induce antiviral gene activation in response to LPS^29, 30^. We have observed strong suppression of the antiviral gene program in response to LPS in PBMCs of opioid-dependent individuals, and very modest suppression of the inflammatory modules (Figure 2). Although there is some evidence of *TLR4* expression in other immune cell types such as CD4+ T cells^31^ and NK cells^32^ in naive PBMCs, monocytes are the major cell type that express high levels of *TLR4* while other immune cell types demonstrate low expression levels as shown from reanalysis of previously published population level RNA-seq data from PBMCs of healthy individuals^28^ (Supplementary Figure S19). We anticipate that LPS induction of the three innate immune response gene pathways by *TLR4* activation occurs mainly in monocytes; this leads to the expression of autocrine and paracrine factors such as *TNFɑ* and *IFNɑ*/*IFNβ* which then induce expression of the innate immune response gene modules in other immune cell types through the activation of TNF receptors and IFN*ɑ*/*β* receptors as shown by the expression of these receptors in CD4+ T cells, CD8+ T cells, B cells and NK cells. In contrast to healthy controls, IFN*β* expression (*IFNB1*) was almost completely suppressed in naive monocytes from opioid-dependent individuals (Supplementary Figure S20). This is consistent with our observation that naive monocytes is the only naive immune cell type that showed suppression of interferon-stimulated genes and antiviral genes (Figure 1).

Furthermore, our study demonstrates the utility of scRNA-seq as an unbiased tool to assess cell type specific genome-wide transcriptomic phenotype from limited quantity of patient samples. Upon a stimulation condition, such as LPS treatment, the opioid induced phenotype is much more pronounced and resembles the signal amplifying effect from electronic amplifiers. We anticipate this type of signal amplification method coupled with single cell transcriptomics will be of broad interest and can be applied to many other disease models where disease relevant stimuli can be used to activate naïve PBMCs isolated from patients that are otherwise quiescent to amplify signal over noisy background and thus reveal the phenotype of the disease.

## Materials and Methods

### PBMC from Opioid-Dependent Individuals

Frozen vials of PBMC prepared from the fresh blood of opioid-dependent (mostly heroin dependent), and non-dependent neighborhood control individuals were collected in the Comorbidity and Trauma Study (CATS)^33, 34^ and subsequently obtained from the biorepository of National Institute on Drug Abuse (NIDA, Rockville MD). We evaluated 14 age-matched subjects ranging from 24-45 years in age, with an equal number of male and female subjects (Supplementary Table 5). Cases were recruited from opioid replacement therapy (ORT) clinics in the greater Sydney, Australia region, while controls were recruited from areas in close proximity to ORT clinics (neighborhood controls). Cases and controls were required to be English speakers 18 years of age or older. Cases were participants in ORT for opioid-dependence while controls were excluded for recreational opioid use 11 or more times lifetime. All subjects provided written informed consent ^33, 34^. All samples were stripped of personally identifying information and assigned sample ID numbers prior to receipt.

### Single Cell Transcriptomic Analysis of PBMC from Opioid-Dependent Individuals

Frozen PBMCs isolated from the blood of opioid-dependent and non-dependent neighborhood individuals were revived, and live cells were isolated via FACS (Fluorescently Activated Cell Sorting) using a Sony SH800 cell sorter and a live/dead cell stain (LIVE/DEAD Fixable Green Cell Stain Kit, for 488nm Excitation, Thermo Fisher – L34969). Dilutions were made from all 14 samples as outlined in the Chromium Single Cell 3’ Reagent Kit v2 User Guide (10X Genomics, CG00052 Rev.B), and 7000 cells per sample were used to perform the droplet-based Chromium Single Cell 3’ scRNA-seq method (10X Genomics, Chromium Single Cell 3’ Library and Gel Bead Kit, Cat# PN-120237). 200,000 cells from 6 of the 14 samples (3 dependent and 3 non-dependent) were plated into a non-tissue culture treated 96-well plate in a leukocyte-supporting complete RPMI medium (10% HI-FBS, 1% L-Glutamate, 1% NEAA, 1% HEPES, 1% Sodium Pyruvate, 0.1% B-Mercaptoethanol). Lipopolysaccharide (LPS) (Invivogen, LPS-EK Ultrapure, Cat# tlrl-pekpls) was then added to a final concentration of 100ng/mL and the cells were incubated at 37°C for 3 hours. Cells were then collected, washed, and diluted to 1000 cells/μL before being used to perform the 10X Genomics Chromium Single Cell 3’ method as outlined in the Single Cell 3’ Reagent Kit v2 User Guide. Briefly, 20μL of 1000 cells/μL PBMC suspension from each subject/condition were combined, and 33.8 μL of cell suspension (total cell number = 33,800) was mixed with 66.2 μL of RT reaction mix before being added to a Chromium microfluidics chip already loaded with barcoded beads and partitioning oil. The chip was then placed within the Chromium controller where single cells and barcoded beads were encapsulated together within oil droplets. Reverse transcription was then performed within the oil droplets to produce barcoded cDNA. Barcoded cDNA was isolated from the partitioning oil using Silane DynaBeads (Thermo Fisher Scientific, Dynabeads MyONE Silane, Cat# 37002D) before amplification by PCR. Cleanup/size selection was performed on amplified cDNA using SPRIselect beads (Beckman-Coulter, SPRIselect, Cat# B23317) and cDNA quality was assessed using an Agilent 2100 BioAnalyzer and the High-Sensitivity DNA assay (Agilent, High-Sensitivity DNA Kit, Cat# 5067-4626). Sequencing Libraries were generated from cleaned, amplified cDNA using the 10X Chromium Kit’s included reagents for fragmentation, sequencing adaptor ligation, and sample index PCR. Between each of these steps, libraries were cleaned/size selected using SPRIselect beads. Final quality of cDNA libraries was once again assessed using the Agilent BioAnalyzer High-Sensitivity DNA assay, and quality-confirmed libraries were sequenced using Illumina’s NextSeq platform. All reagents listed in Supplementary Table 2.

### PBMC from healthy Individuals used in In vitro assays

15 mL of fresh whole blood from healthy donors (Research Blood Components, Boston MA) was diluted 1:1 with warm PBS +2% FBS, mixed, and gently layered atop 30mL Ficoll-Paque density gradient medium (GE Healthcare, Ficoll Paque PLUS, Cat# 17-1440) in 50mL conical tubes. This process was repeated 7 times in processing 100mL of blood. Tubes were centrifuged for 20 min at 1200xg to separate leukocytes from red blood cells and plasma. The leukocyte-containing buffy coat was carefully transferred into new tubes, washed with warm PBS +2% FBS, counted, resuspended in DMSO and aliquoted. Isolated cells were then stored in liquid nitrogen until later experimental use. All reagents listed in Supplementary Table 2.

### Controlled Substances

Solid Morphine Sulfate (Sigma Aldritch, Cat# M8777-25G) was obtained with approval and oversight from the controlled substances sub-office of the Boston University department of environmental health and safety. Aliquots of a 10mM stock solution were prepared and stored for further use in experimentation. All reagents listed in Supplementary Table 2

### Morphine Titration in PBMCs from healthy individuals

Normal PBMC were revived in leukocyte-supporting complete RPMI medium and plated onto non-tissue culture treated 96 well plates at a density of 2.0e5 cells/well (2 wells per condition, 4.0e5 cells total). Cells were treated either with a mock treatment or a titration of Morphine Sulfate in RPMI complete medium (0μM, 10μM, 100μM) for 24 hours. At the end of the morphine incubation either medium or LPS (final concentration 100ng/mL) was added to the wells and cells were incubated for a further 3 hours, at the end of which the cells were collected, washed, and processed for total RNA using the ZymoPure QuickRNA MiniPrep kit (Zymo Research, Cat# R1055). RNA samples were then used to perform RT-qPCR. All reagents listed in Supplementary Table 2

### RT-qPCR analysis

Total RNA was isolated from cells using the ZymoPure QuickRNA MiniPrep kit. cDNA was synthesized using ∼50ng of total RNA per sample (Thermo Fisher, SuperScript IV First-Strand Synthesis System, Cat# 18091200). 2 μL of cDNA per reaction was then used to perform qPCR (Fisher Scientific, PowerUp SYBR Greβen Master Mix, Cat# A25742) with primers against transcripts of the Interferon target gene ISG15 (IDT, primer sequences in supplement) We used primers against ActB (IDT, primer sequences in supplement) as a housekeeping gene control. All reagents listed in Supplementary Table 2.

### Cell Hashing scRNA-seq

Morphine titration assays were performed as previously described. At the end of the treatment period cells were collected, washed, and each sample was “hashed” using unique oligonucleotide-barcoded antibodies ^16^ (Supplementary Table 3) to track the cells’ well/condition of origin. Briefly, cells were suspended in Cell Staining Buffer (BioLegend, Cat# 420201) and blocked using Human TruStain FcX reagent (BioLegend, Cat# 422301). Cells were then incubated with TotalSeq antibodies (BioLegend, Cat# 3964601, 394603, 394605, 394607, 394609, 394611, 394613, 394615, 394617, 394619, 394623, 394625), washed with PBS, and filtered through 40μM cell strainers (Bel-Art, Flowmi Cell Strainer, Cat# H13680-0040). Samples were then normalized to 1000 cells/μL, mixed in equal measure (20μL each), and used to perform the Chromium Single Cell 3’ scRNA-seq method as described previously. Additional primers were included in the cDNA amplification step to account for the TotalSeq oligonucleotide tags. During the post-amplification cleanup, supernatant containing amplified TotalSeq tags was collected and processed parallel to the standard 10X library fraction. All reagents listed in Supplementary Table 2.

### Single Cell Analysis

#### RNA-sequencing preprocessing

We used Cell Ranger version 2.1.0 (10x Genomics) to pool and process the raw RNA sequencing data. First, each sample sequencing library was demultiplexed based on the sample index read to generate FASTQ files for the paired-end reads. STAR aligner^35^ was used to align reads to the human reference genome (GRCh38). After alignment, barcode and UMI filtering as well as UMI counting were performed to then generate the gene-cell barcode matrix for each sample library.

To perform an integrative analysis with the opioid-dependent and non-dependent control samples, all samples were normalized by sequencing depth and each sample gene-cell barcode matrix was concatenated together to create a large gene-barcode matrix in which cell-to-cell and sample-to-sample comparisons could be made.

#### RNA-sequencing downstream and DE analysis

We used Seurat suite version 2.3^10, 11^ for downstream analysis. We used Seurat quality control metrics to select genes and cells for further analysis. Cells with less than 300 and greater than 2,000 detected genes were filtered out, as well as cells with greater than 10,000 UMIs were filtered out. Genes that were detected in less than 10 cells were removed. Gene counts for each cell were normalized by total expression, multiplied by a scale factor of 10,000 and transformed to log scale. Principal component analysis based on the highly variable genes detected (dispersion of 2) was performed for dimension reduction and the top 20 principal components (PCs) were selected. We clustered cells based on graph-based methods (KNN and Louvain community detection method) implemented in Seurat. The clusters and other known annotations were visualized using t-stochastic distributed embedding (tSNE)^36^.

To identify peripheral immune subpopulations, we performed differential expression analysis using Wilcoxon rank sum test between clusters to identify top expressing genes for each cluster for cell type identification implemented in Seurat. Cell type specific gene signatures were determined from the overlap of more highly expressed and canonical gene markers. In addition, we performed differential expression analysis within each cell type between control cells and opioid-dependent cells using Model-based Analysis of Single Cell Transcriptomics (MAST)^37^ to identify genes upregulated and downregulated in opioid-dependent patients compared to controls (Supplementary Table 4). MAST uses a Hurdle model, a two-part generalized linear model in which one process accounts for zero expression and the second process accounts for positive expression. Differentially expressed genes were evaluated according to their log fold change (greater than log2(1.5)) and adjusted p-values (0.05). All statistical tests in the analysis of expression of the three innate immune response gene modules activated through LPS stimulation^12^ (Supplementary Table 1) between opioid-dependent cells and control cells were performed using t-tests through the ggpubr R package (https://github.com/kassambara/ggpubr).

#### Cell Hashing preprocessing and analysis

For hashtag oligo (HTO) quantification, we first ran Cite-seq-Count^16, 38^ on the HTO fastq files to process the HTO reads with the parameters specific to 10x Genomics single cell 3’ v2 data as stated in (https://github.com/Hoohm/CITE-seq-Count). In addition, we used Cell Ranger version 2.1.1 (10x Genomics) to process the raw sequencing RNA reads and Seurat suite version 2.3 for downstream analyses. To identify the cells sample-of-origin, we demultiplexed the HTOs and removed doublets and ambiguous cells using the Seurat pipeline for demultiplexing as mentioned in (https://satijalab.org/seurat/hashing_vignette.html).

#### Enrichment Analysis

We performed gene enrichment analysis of the list of differential genes between opioid-dependent individuals and non-dependent neighborhood controls for each cell type using Metascape^39^ online tool (http://metascape.org/). The enrichment analysis was run using default settings, and was assessed and visualized through a heatmap of significance (-log(P-value)). All heatmaps were generated using ComplexHeatmap R package^40^.

## Acknowledgements

We thank Dr. Todd A. Blute for his technical support on flow cytometry analysis. We would like to thank Dr. David J. Waxman and Dr. Andrew J. Henderson for critical reading of the manuscript. The work was supported by NIH grant R61DA047032. T.T.K was supported by NIH institutional training grant T32GM100842. CATS data collection was funded by R01DA17305 and E.C.N. is supported by R01 DA042620, R01 DA046436, and R33 DA041883.

## Data Availability

Processed cell hashing scRNA-seq data is available from GEO under accession GSE128879. Raw and processed scRNA-seq data of opioid-dependent individuals and neighborhood controls is available from dbGaP under accession phs000277.v2.p1.

---

**Figure S1.**
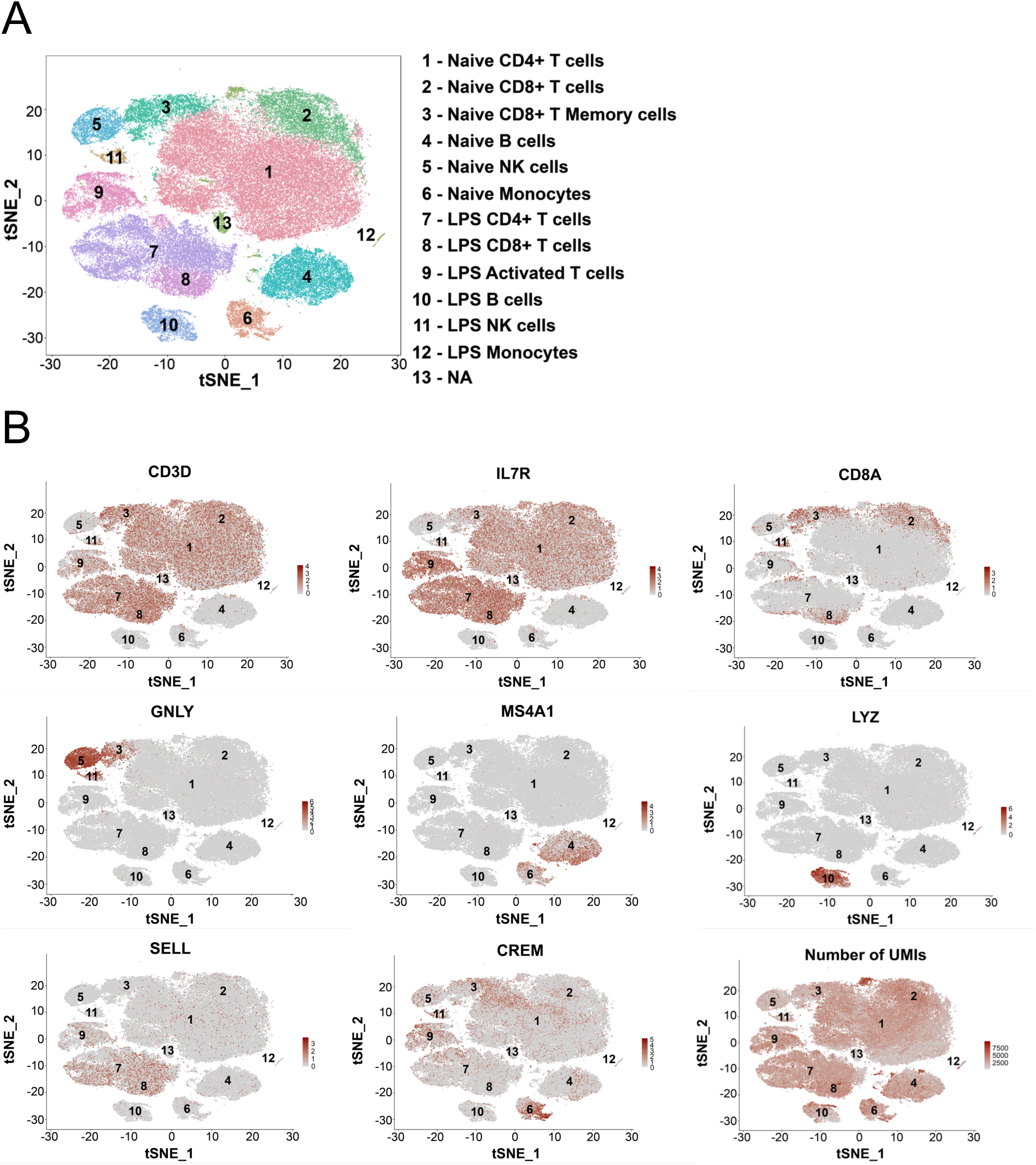
Cell type identification from scRNA-seq of naive and LPS-stimulated PBMCs from opioid-dependent individuals and neighborhood controls using canonical gene markers. **a,** t-SNE plot of naive and LPS-stimulated PBMCs with identified cell types: Naive CD4+ T cells (30,388 cells), Naive CD8+ T cells (6,439 cells), Naive CD8+ memory T cells (3,588 cells), Naive B cells (6,703 cells), Naive NK cells (2,439 cells), Naive monocytes (2,220 cells), LPS-stimulated CD4+ T cells (11,308 cells), LPS-stimulated CD8+ T cells (3,019 cells), LPS-stimulated activated T cells (4,058 cells), LPS-stimulated B cells (2,225 cells), LPS-stimulated NK cells (566 cells), LPS-stimulated monocytes (261 cells), NA or no applicable cell types (1,608 cells). **b,** t-SNE projection of canonical gene marker expression and number of UMIs per cell across all naive state and LPS-stimulated subpopulations: CD4+ T cells (*CD3D*, *IL7R*), CD8+ T cells (*CD8A*), B cells (*MS4A1*), NK cells (*GNLY*), Monocytes (*LYZ*), activated cells (*CREM*), naive cells (*SELL*).

**Figure S2.**
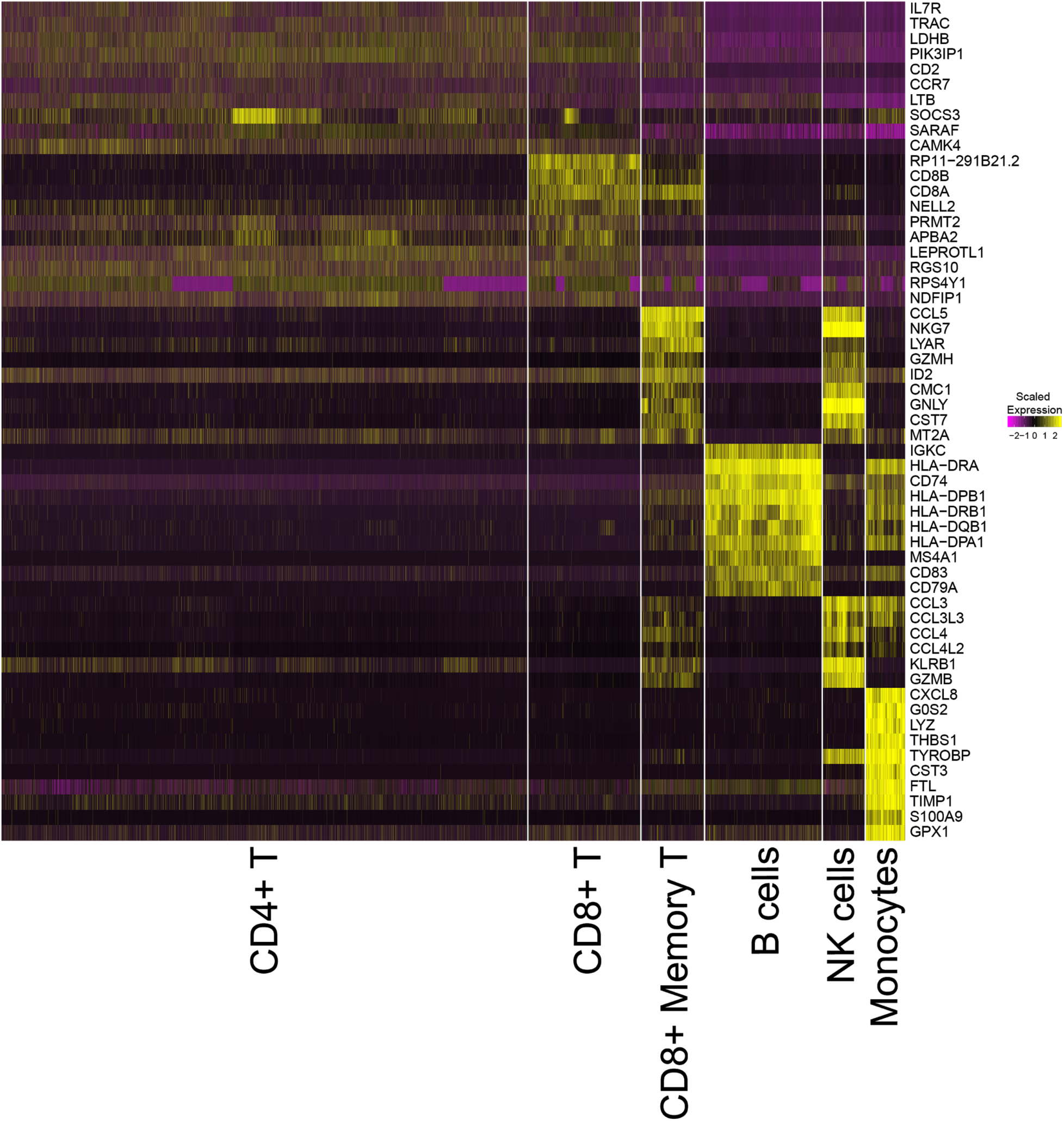
Top marker genes for naive state cell types. Heatmap of the top 10 genes expressed (scaled) across cells in each naive state population. Top 10 marker genes were identified using a wilcoxon rank sum test over all cell types: CD4+ T cells (CD4+ T), CD8+ T cells (CD8+ T), CD8+ memory T cells (CD8+ T Memory), B cells, NK cells, and Monocytes.

**Figure S3.**
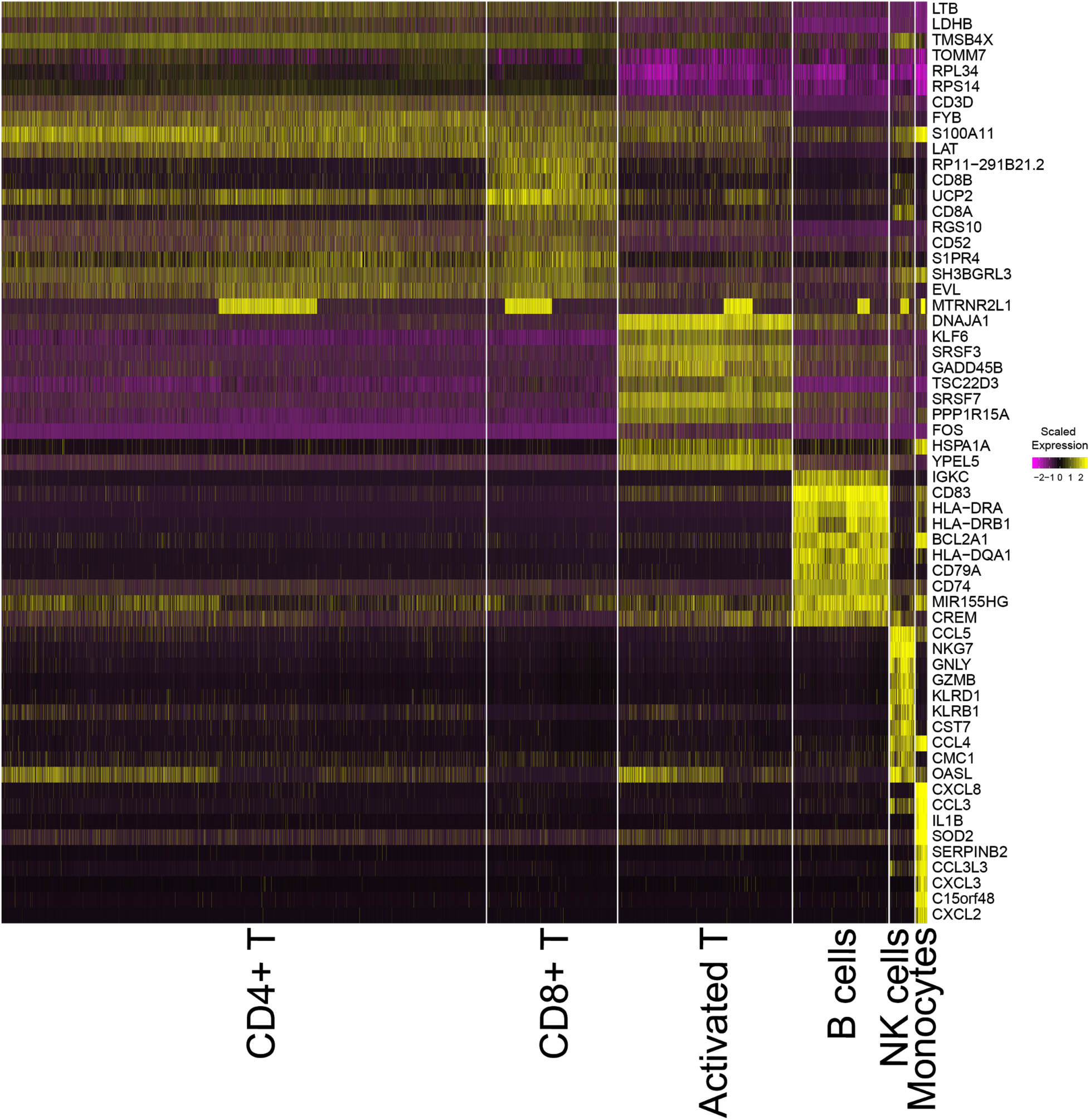
Top marker genes for LPS-stimulated state cell types. Heatmap of the top 10 gene markers expressed (scaled) across cells in each LPS-stimulated population. Top 10 marker genes were identified using a wilcoxon rank sum test over all cell types: CD4+ T cells (CD4+ T), CD8+ T cells (CD8+ T), activated T cells (Activated T), B cells, NK cells, and Monocytes.

**Figure S4.**
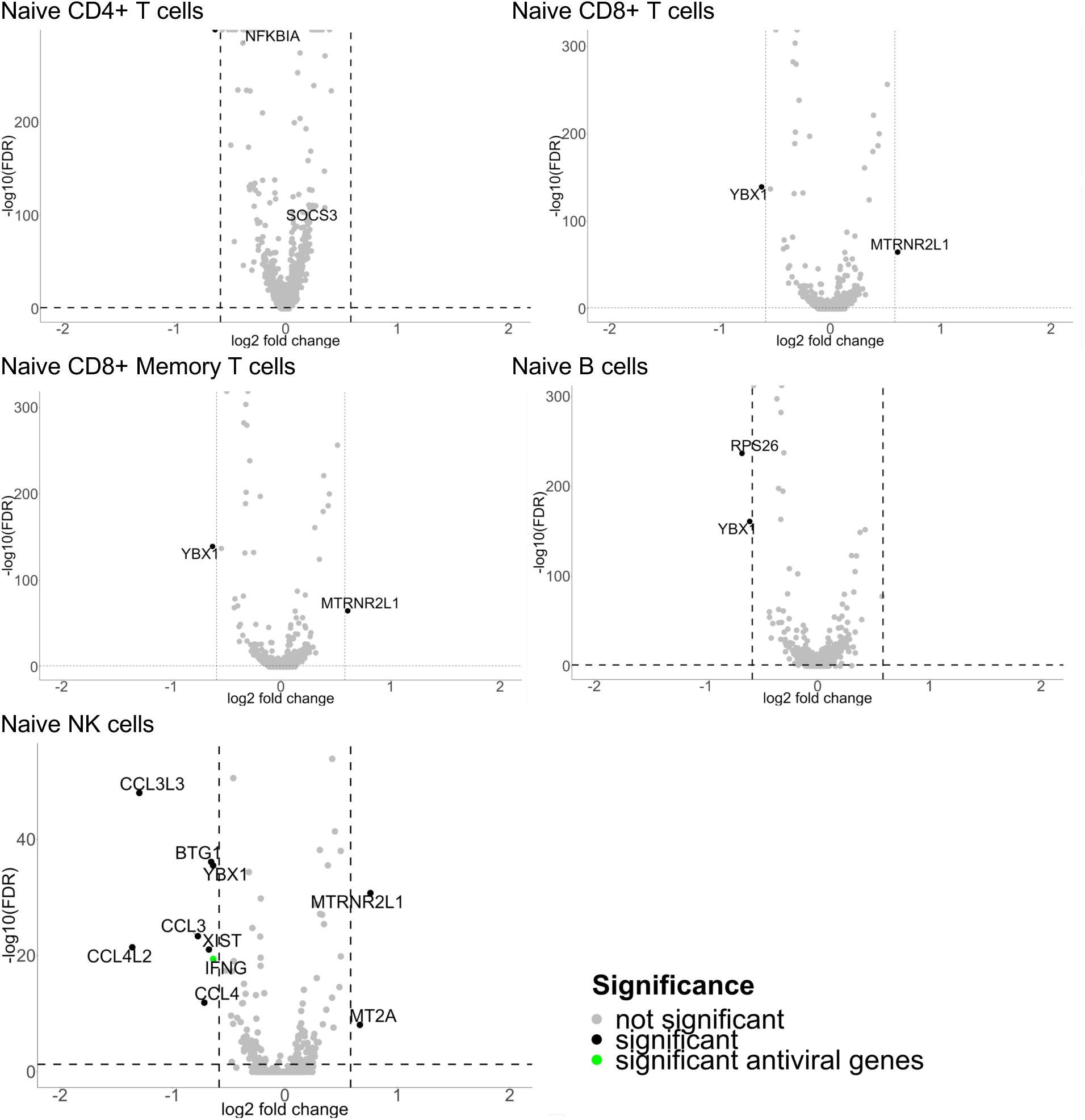
Differential gene expression analysis of opioid-dependent and neighborhood control PBMCs across naive state cell types. We performed differential expression analysis within each cell type between control and opioid-dependent cells (see Methods). Volcano plot showing fold change of genes (log2 scale) for opioid-dependent cells compared to controls from Naive CD4+ T cells, Naive CD8+ T cells, Naive CD8+ memory T cells, Naive B cells, and Naive NK cells (x-axis) and significance of 0.05 (y-axis, −log10 scale). Significant genes shown in black, significant antiviral genes shown in green, and insignificant genes shown in grey.

**Figure S5.**
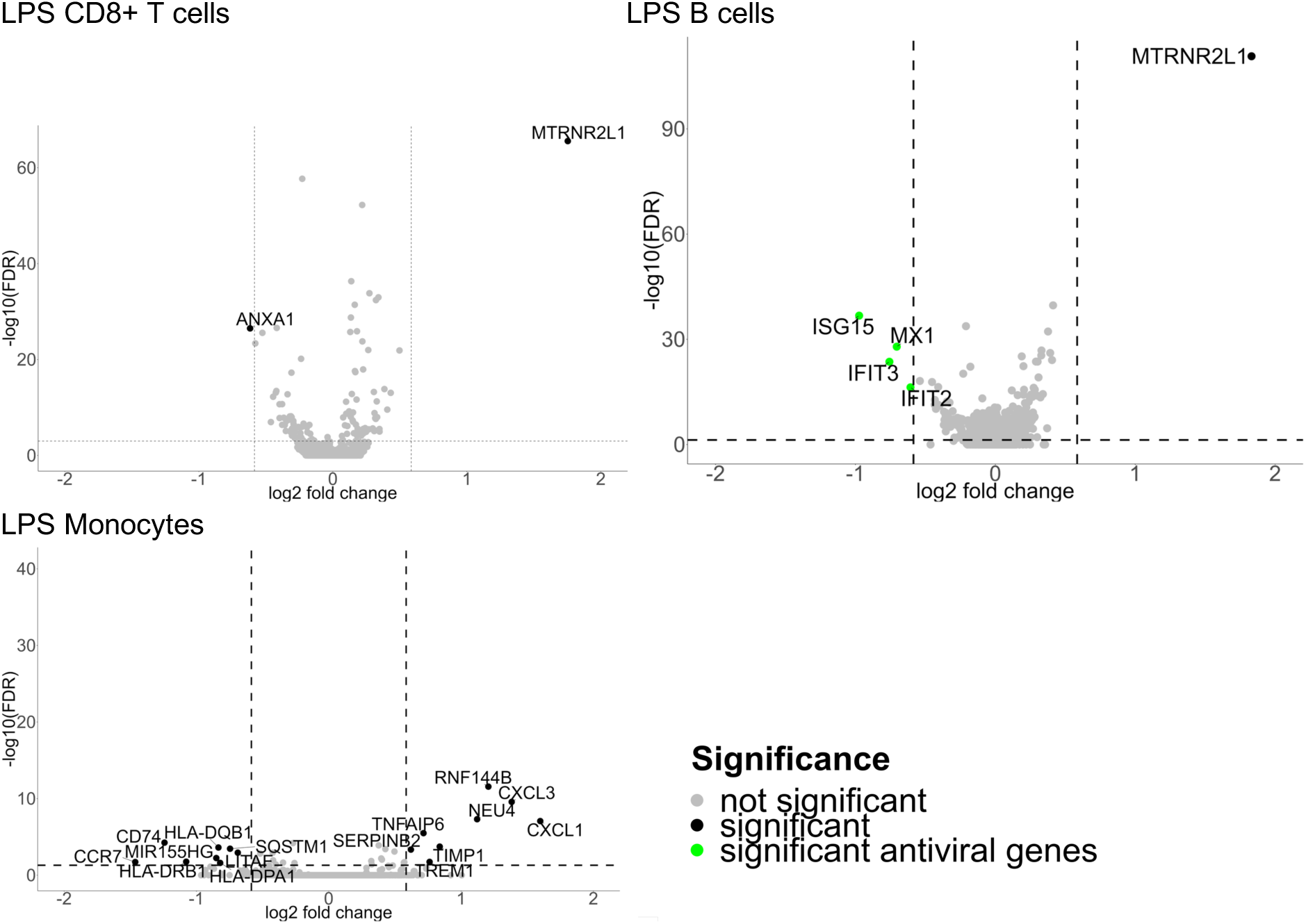
Differential gene expression analysis of opioid-dependent and neighborhood control PBMCs across LPS-stimulated state cell types. We performed differential expression analysis within each cell type between control and opioid dependent cells (see Methods). Volcano plot showing fold change of genes (log2 scale) for opioid-dependent cells compared to controls from LPS-stimulated CD8+ T cells, LPS-stimulated B cells, LPS-stimulated Monocytes (x-axis) and significance of 0.05 (y-axis, −log10 scale). Significant genes shown in black, significant antiviral genes shown in green, and insignificant genes shown in grey.

**Figure S6.**
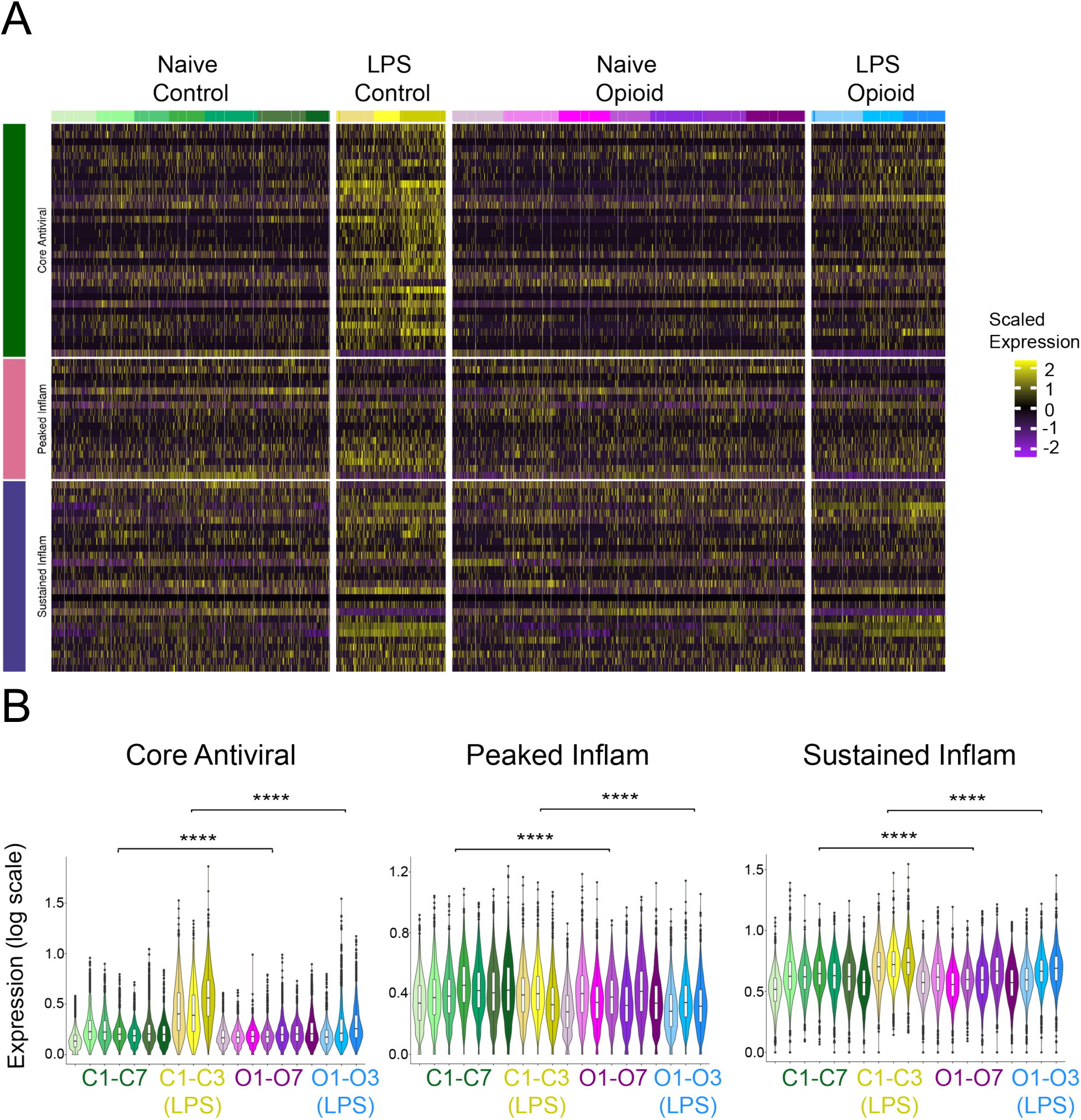
Single cell gene expression heatmap of antiviral and inflammatory gene modules in CD4+ T cells. **a,** Heatmap of scaled expression of core antiviral, peaked inflammatory, and sustained inflammatory gene modules (y-axis) for samples cells: naive state control samples (Naive Control), LPS-stimulated control samples (LPS Control), naive state opioid-dependent samples (Naive Opioid), and LPS-stimulated opioid-dependent samples (LPS Opioid). **b,** Average antiviral gene set expression (log expression) across all sample cells in CD4+ T cells: C1-C7 (naive control samples 1 - 7), C1-C3 (LPS) (LPS-stimulated control samples 1 - 3), O1-O7 (naive opioid dependent samples 1-7), O1-O3 (LPS) (LPS-stimulated opioid dependent samples 1-3). Significance shown using p-values: * < 0.05, ** < 0.01, *** < 0.001, **** < 0.0001.

**Figure S7.**
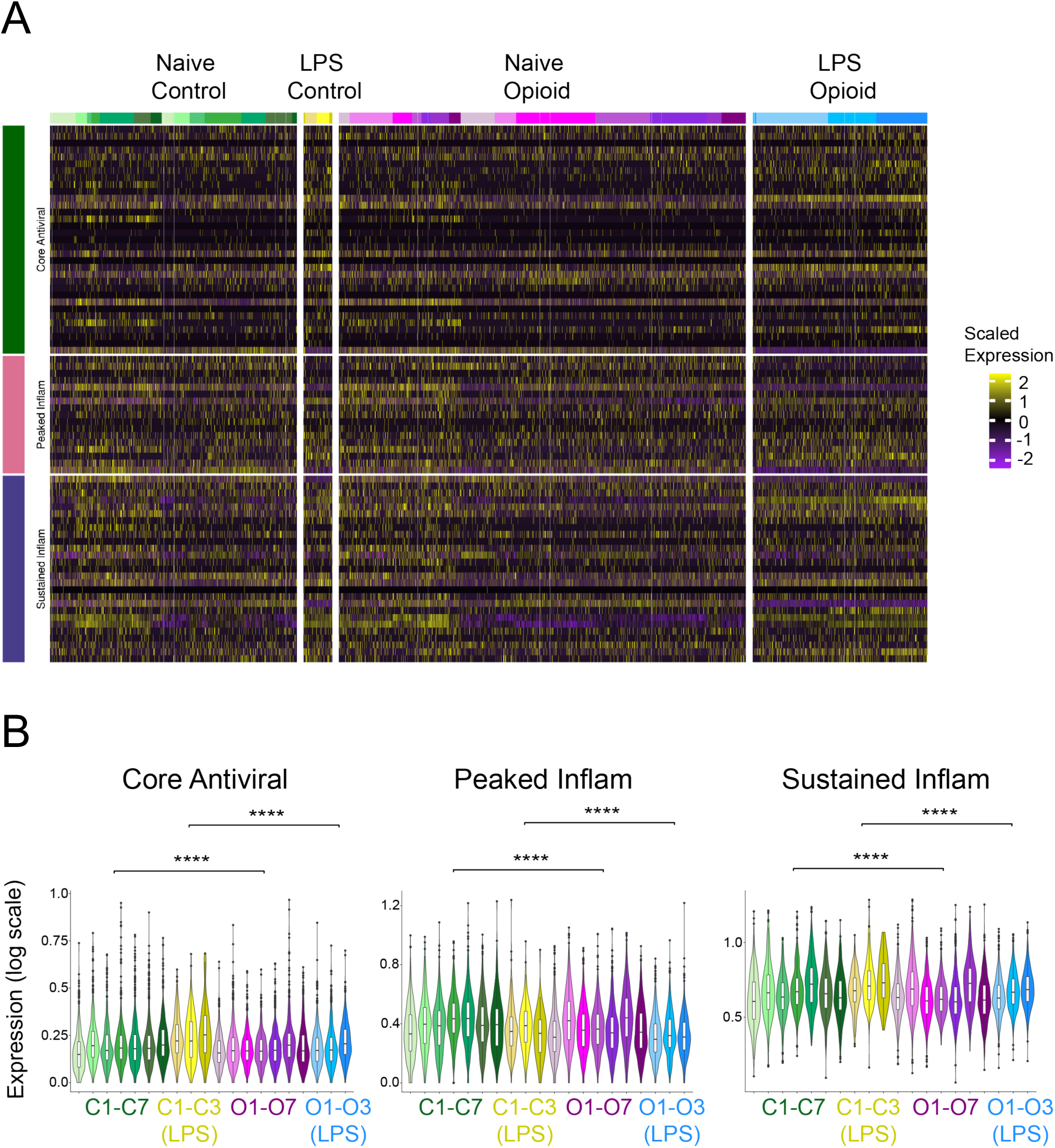
Single cell gene expression heatmap of antiviral and inflammatory gene modules in CD8+ T cells. **a,** Heatmap of scaled expression of core antiviral, peaked inflammatory, and sustained inflammatory gene modules (y-axis) for samples cells: naive state control samples (Naive Control), LPS-stimulated control samples (LPS Control), naive state opioid-dependent samples (Naive Opioid), and LPS-stimulated opioid-dependent samples (LPS Opioid). **b,** Average antiviral gene set expression (log expression) across all sample cells in CD8+ T cells: C1-C7 (naive control samples 1 - 7), C1-C3 (LPS) (LPS-stimulated control samples 1 - 3), O1-O7 (naive opioid dependent samples 1-7), O1-O3 (LPS) (LPS-stimulated opioid dependent samples 1-3). Significance shown using p-values: * < 0.05, ** < 0.01, *** < 0.001, **** < 0.0001.

**Figure S8.**
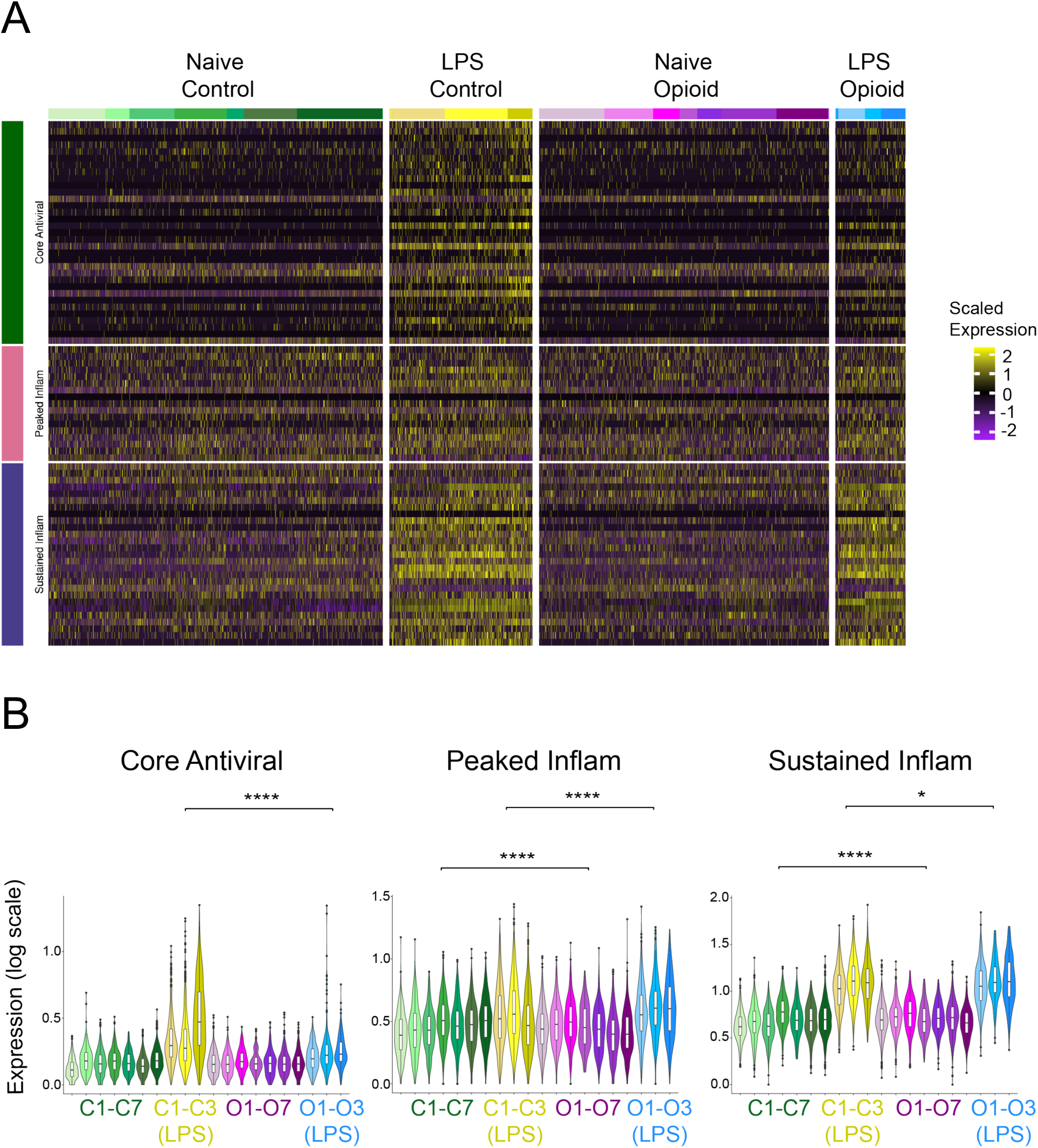
Single cell gene expression heatmap of antiviral and inflammatory gene modules in B cells. **a,** Heatmap of scaled expression of core antiviral, peaked inflammatory, and sustained inflammatory gene modules (y-axis) for samples cells: naive state control samples (Naive Control), LPS-stimulated control samples (LPS Control), naive state opioid-dependent samples (Naive Opioid), and LPS-stimulated opioid-dependent samples (LPS Opioid). **b,** Average antiviral gene set expression (log expression) across all sample cells in B cells: C1-C7 (naive control samples 1 - 7), C1-C3 (LPS) (LPS-stimulated control samples 1 - 3), O1-O7 (naive opioid dependent samples 1-7), O1-O3 (LPS) (LPS-stimulated opioid dependent samples 1-3). Significance shown using p-values: * < 0.05, ** < 0.01, *** < 0.001, **** < 0.0001.

**Figure S9.**
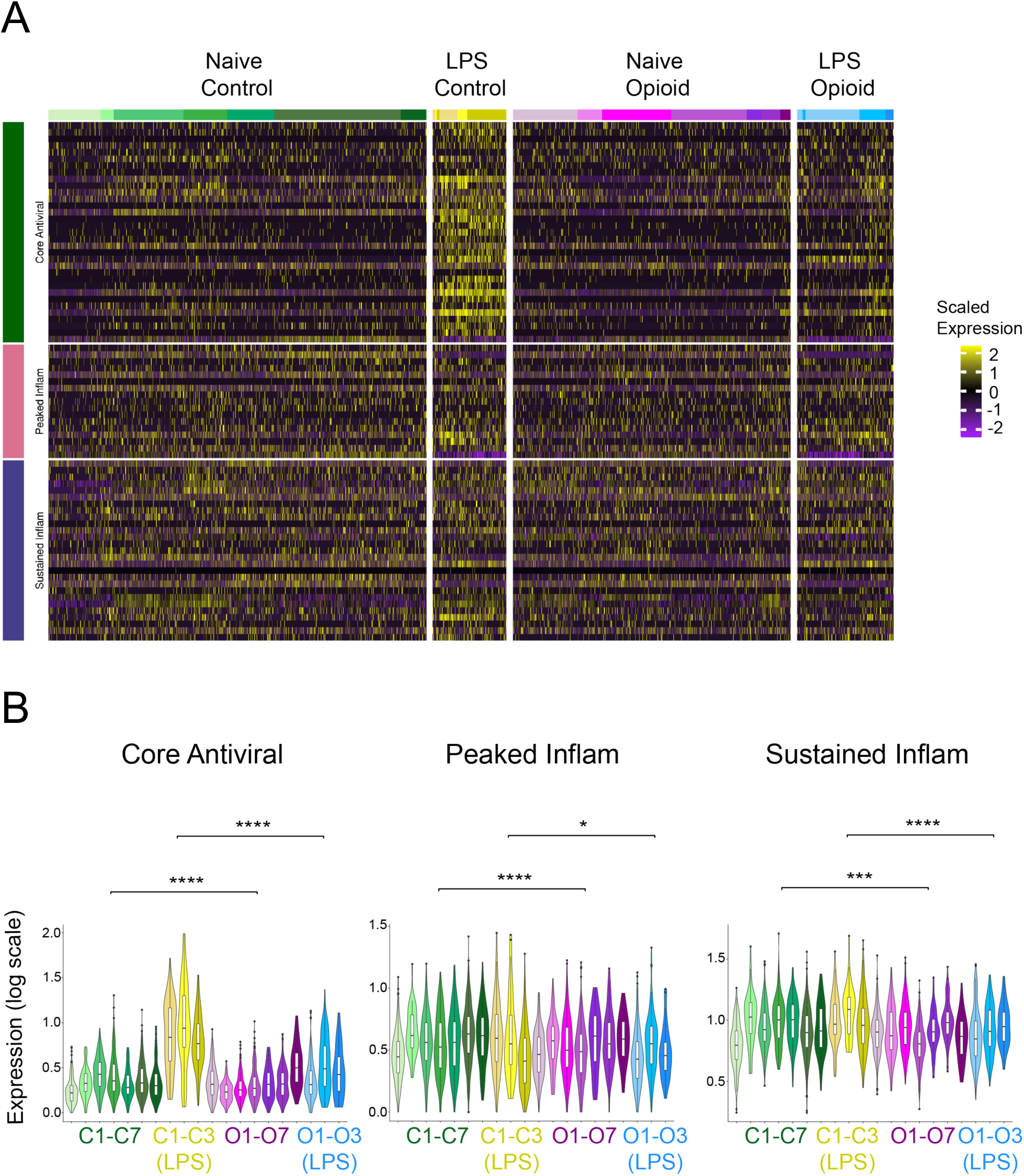
Single cell gene expression heatmap of antiviral and inflammatory gene modules in NK cells. **a,** Heatmap of scaled expression of core antiviral, peaked inflammatory, and sustained inflammatory gene modules (y-axis) for samples cells: naive state control samples (Naive Control), LPS-stimulated control samples (LPS Control), naive state opioid-dependent samples (Naive Opioid), and LPS-stimulated opioid-dependent samples (LPS Opioid). **b,** Average antiviral gene set expression (log expression) across all sample cells in NK cells: C1-C7 (naive control samples 1 - 7), C1-C3 (LPS) (LPS-stimulated control samples 1 - 3), O1-O7 (naive opioid dependent samples 1-7), O1-O3 (LPS) (LPS-stimulated opioid dependent samples 1-3). Significance shown using p-values: * < 0.05, ** < 0.01, *** < 0.001, **** < 0.0001.

**Figure S10.**
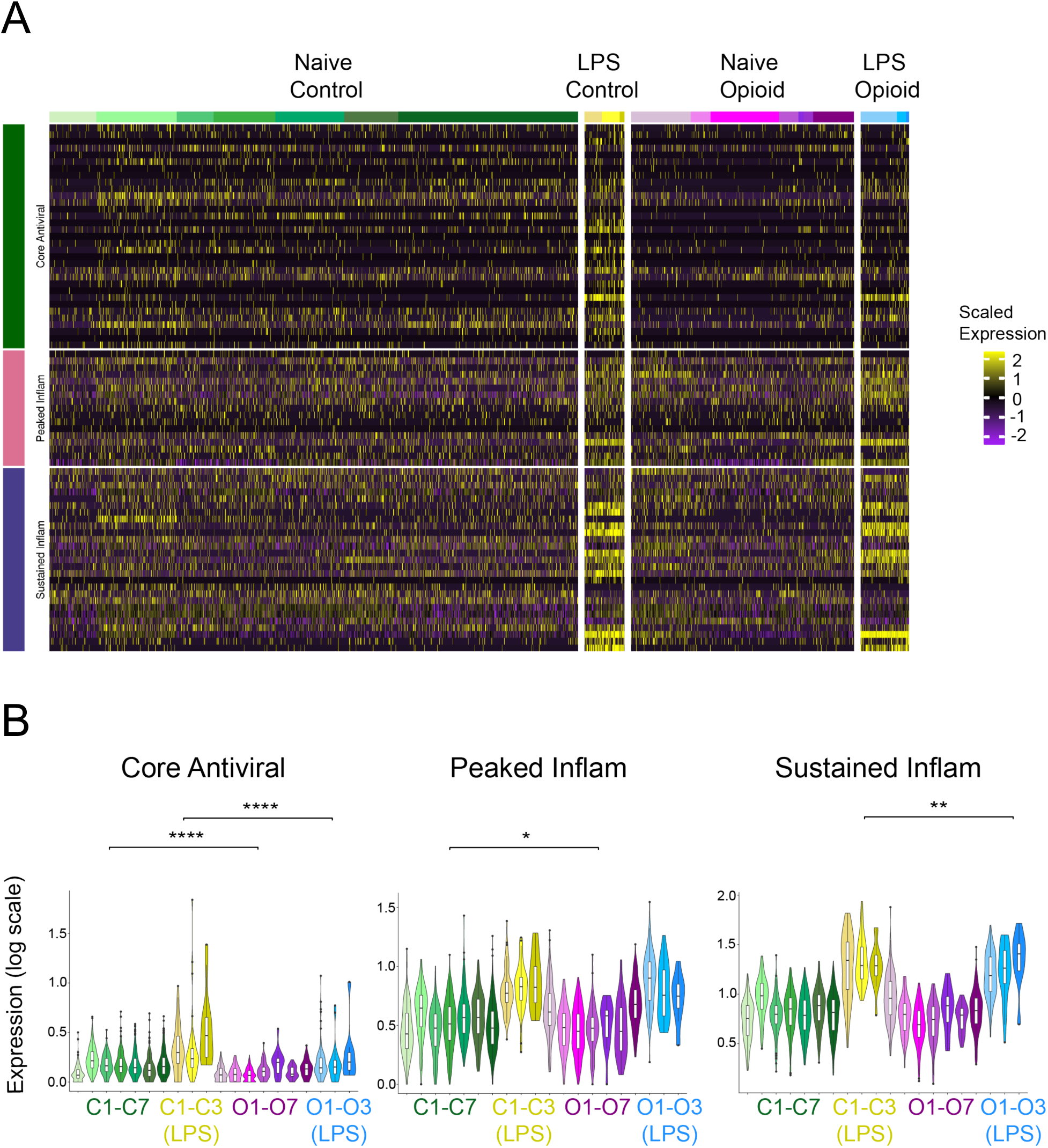
Single cell gene expression heatmap of antiviral and inflammatory gene modules in Monocytes. **a,** Heatmap of scaled expression of core antiviral, peaked inflammatory, and sustained inflammatory gene modules (y-axis) for samples cells: naive state control samples (Naive Control), LPS-stimulated control samples (LPS Control), naive state opioid-dependent samples (Naive Opioid), and LPS-stimulated opioid-dependent samples (LPS Opioid). **b,** Average antiviral gene set expression (log expression) across all sample cells in monocytes: C1-C7 (naive control samples 1 - 7), C1-C3 (LPS) (LPS-stimulated control samples 1 - 3), O1-O7 (naive opioid dependent samples 1-7), O1-O3 (LPS) (LPS-stimulated opioid dependent samples 1-3). Significance shown using p-values: * < 0.05, ** < 0.01, *** < 0.001, **** < 0.0001.

**Figure S11.**
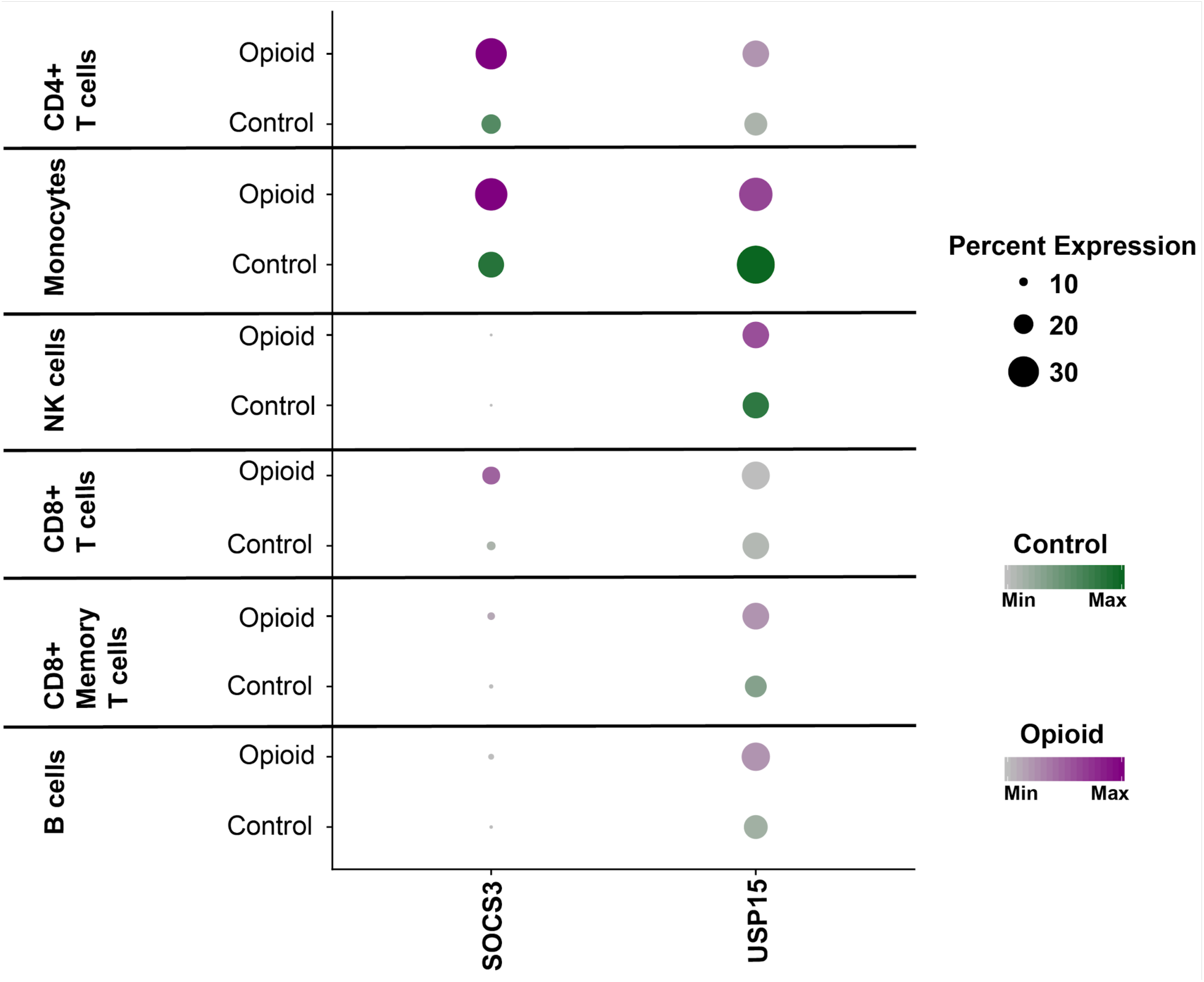
Gene expression of interferon inhibitors in opioid-dependent and control cells. Dot plot of average gene expression of negative interferon regulators *SOCS3* and *USP15* (x-axis) in control cells (Control) and opioid-dependent cells (Opioid) across all naive state cell types: CD4+ T cells, Monocytes, NK cells, CD8+ T cells, CD8+ Memory T cells, and B cells (y-axis).

**Figure S12.**
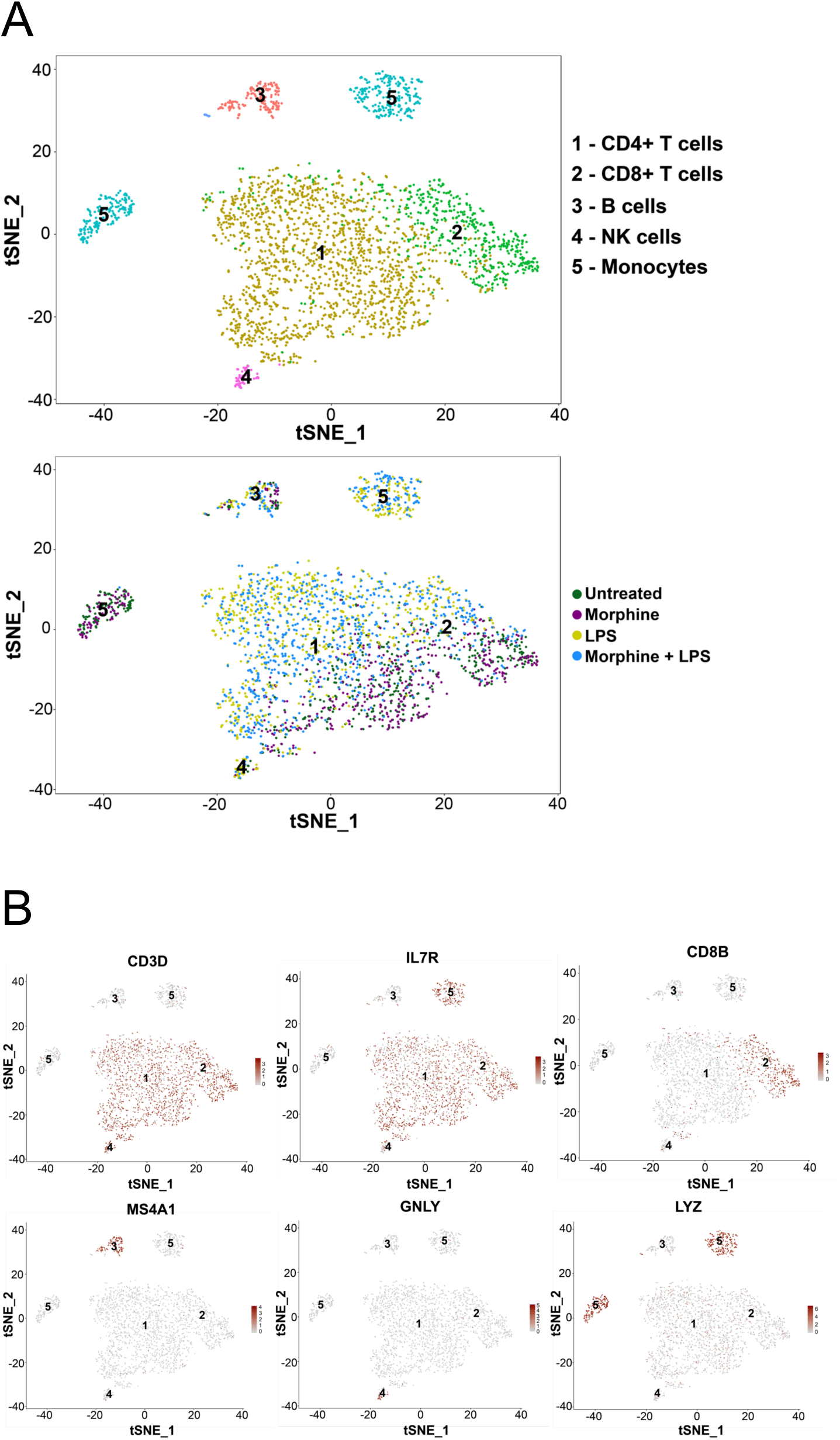
Cell type identification of in vitro morphine-treated PBMCs. **a,** t-SNE projection of unsupervised clustering of 2,958 HTO cells. We identified cell type populations (top): CD4+ T cells (1,774 cells), CD8+ T cells (546 cells), B cells (152 cells), NK cells (58 cells), Monocytes (416 cells). We also colored cells by the 4 HTO samples in naive and LPS-stimulated states (bottom). **b,** t-SNE projection of canonical gene marker expression across all subpopulations: CD4+ T cells (*CD3D*, *IL7R*), CD8+ T cells (*CD3D*, *CD8B*), B cells (*MS4A1*), NK cells (*GNLY*), and Monocytes (*LYZ*).

**Figure S13.**
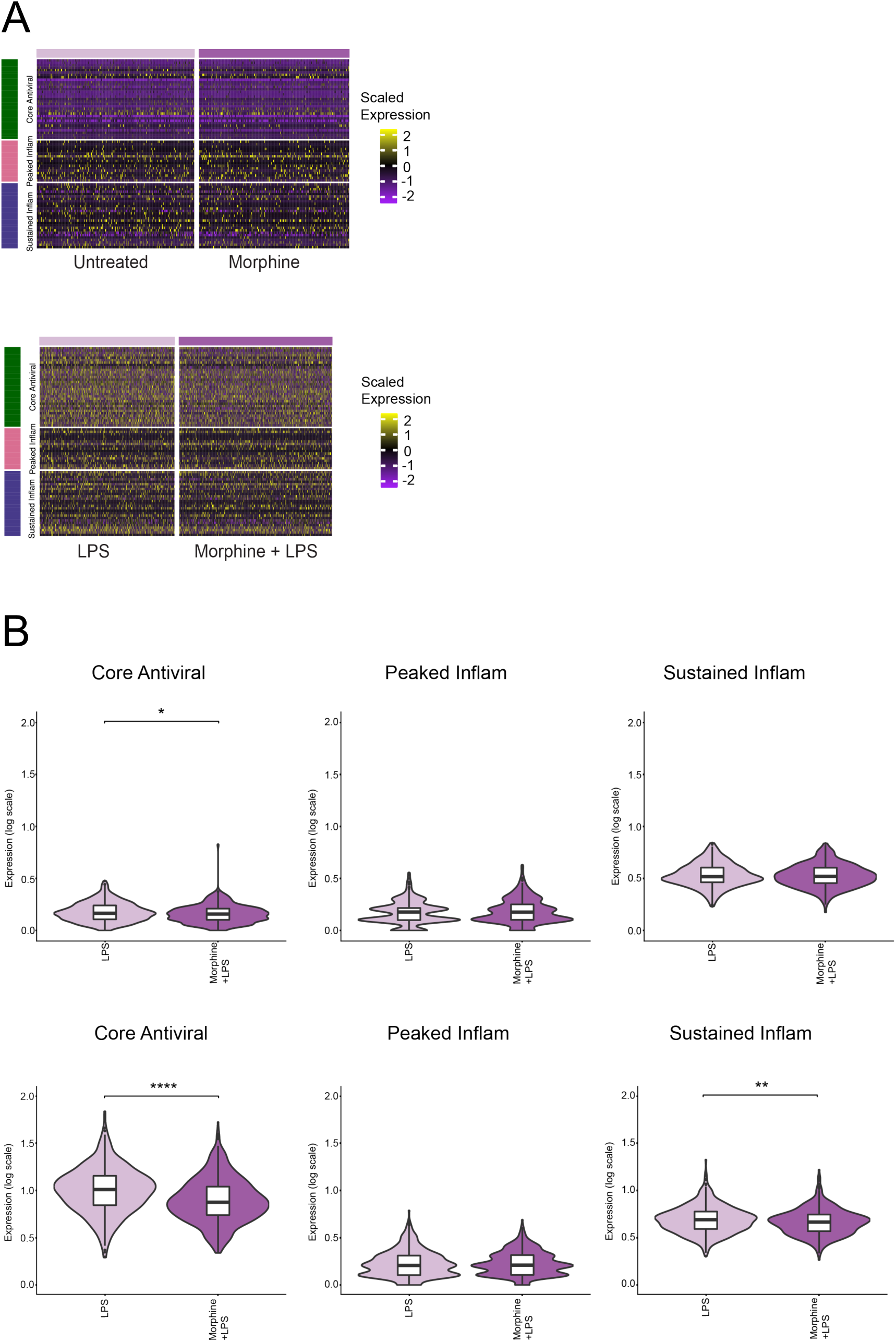
Single cell gene expression heatmap of antiviral and inflammatory gene modules in in vitro treated CD4+ T cells. **a,** Heatmap of scaled expression of core antiviral, peaked inflammatory, and sustained inflammatory gene modules observed across all naive state samples (top) and LPS-stimulated samples (bottom). **b,** Average antiviral gene set expression (log expression) across all naive state sample cells (top) and LPS-stimulated sample cells (bottom). Significance shown using p-values: * < 0.05, ** < 0.01, *** < 0.001, **** < 0.0001.

**Figure S14.**
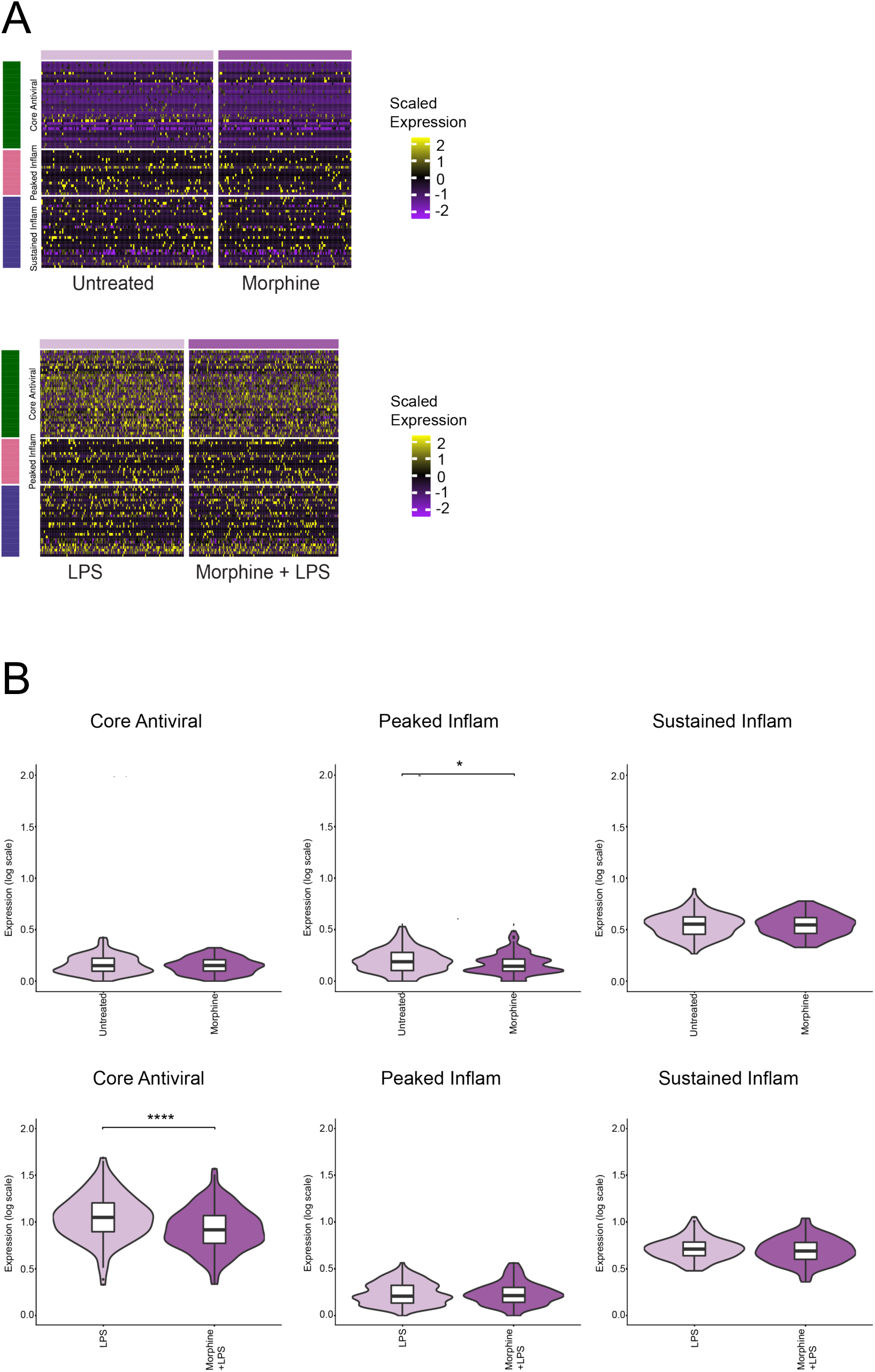
Single cell gene expression heatmap of antiviral and inflammatory gene modules in in vitro treated CD8+ T cells. **a,** Heatmap of scaled expression of core antiviral, peaked inflammatory, and sustained inflammatory gene modules observed across all naive state samples (top) and LPS-stimulated samples (bottom). **b,** Average antiviral gene set expression (log expression) across all naive state sample cells (top) and LPS-stimulated sample cells (bottom). Significance shown using p-values: * < 0.05, ** < 0.01, *** < 0.001, **** < 0.0001.

**Figure S15.**
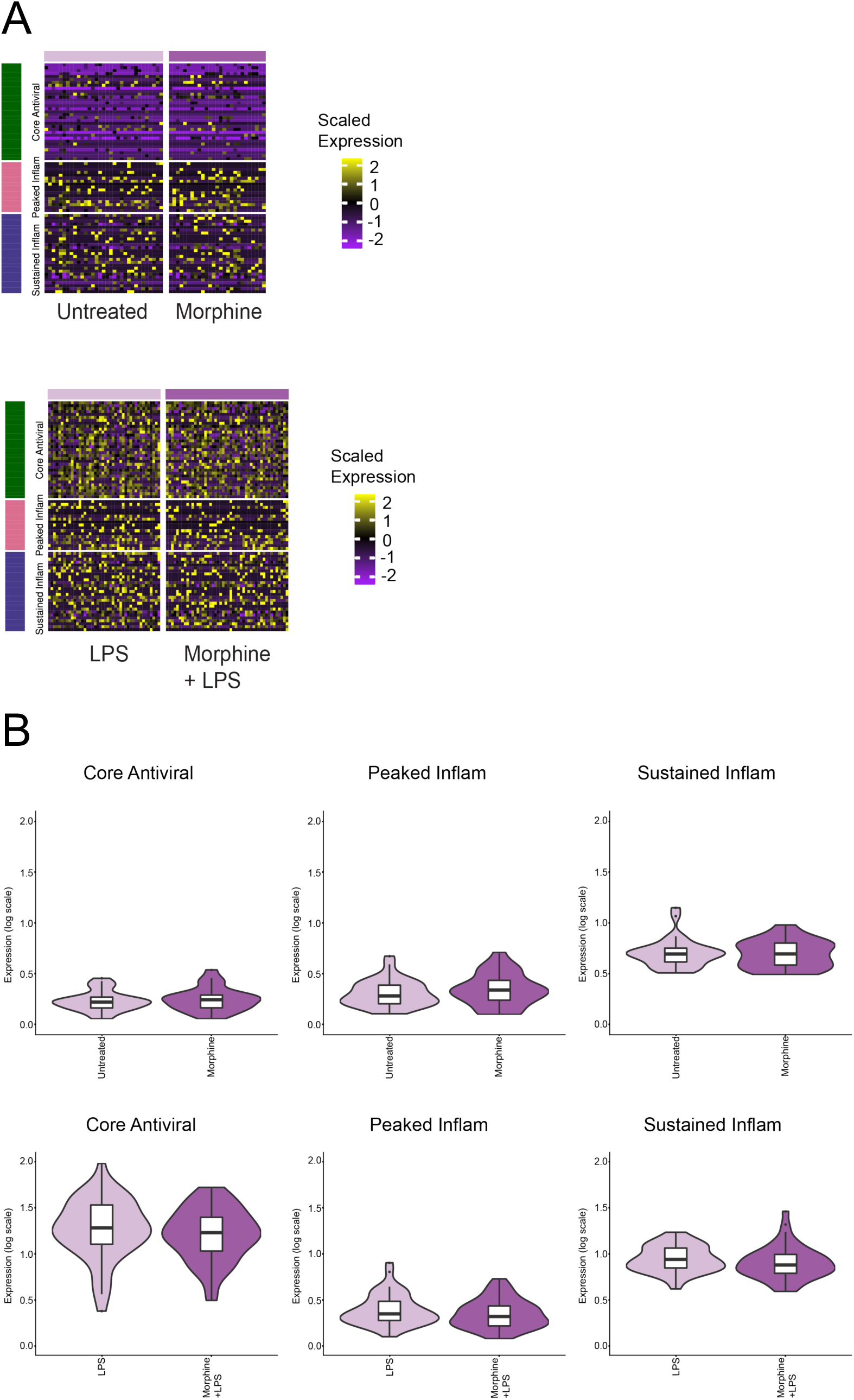
Single cell gene expression heatmap of antiviral and inflammatory gene modules in in vitro treated B cells. **a,** Heatmap of scaled expression of core antiviral, peaked inflammatory, and sustained inflammatory gene modules observed across all naive state samples (top) and LPS-stimulated samples (bottom). **b,** Average antiviral gene set expression (log expression) across all naive state sample cells (top) and LPS-stimulated sample cells (bottom). Significance shown using p-values: * < 0.05, ** < 0.01, *** < 0.001, **** < 0.0001.

**Figure S16.**
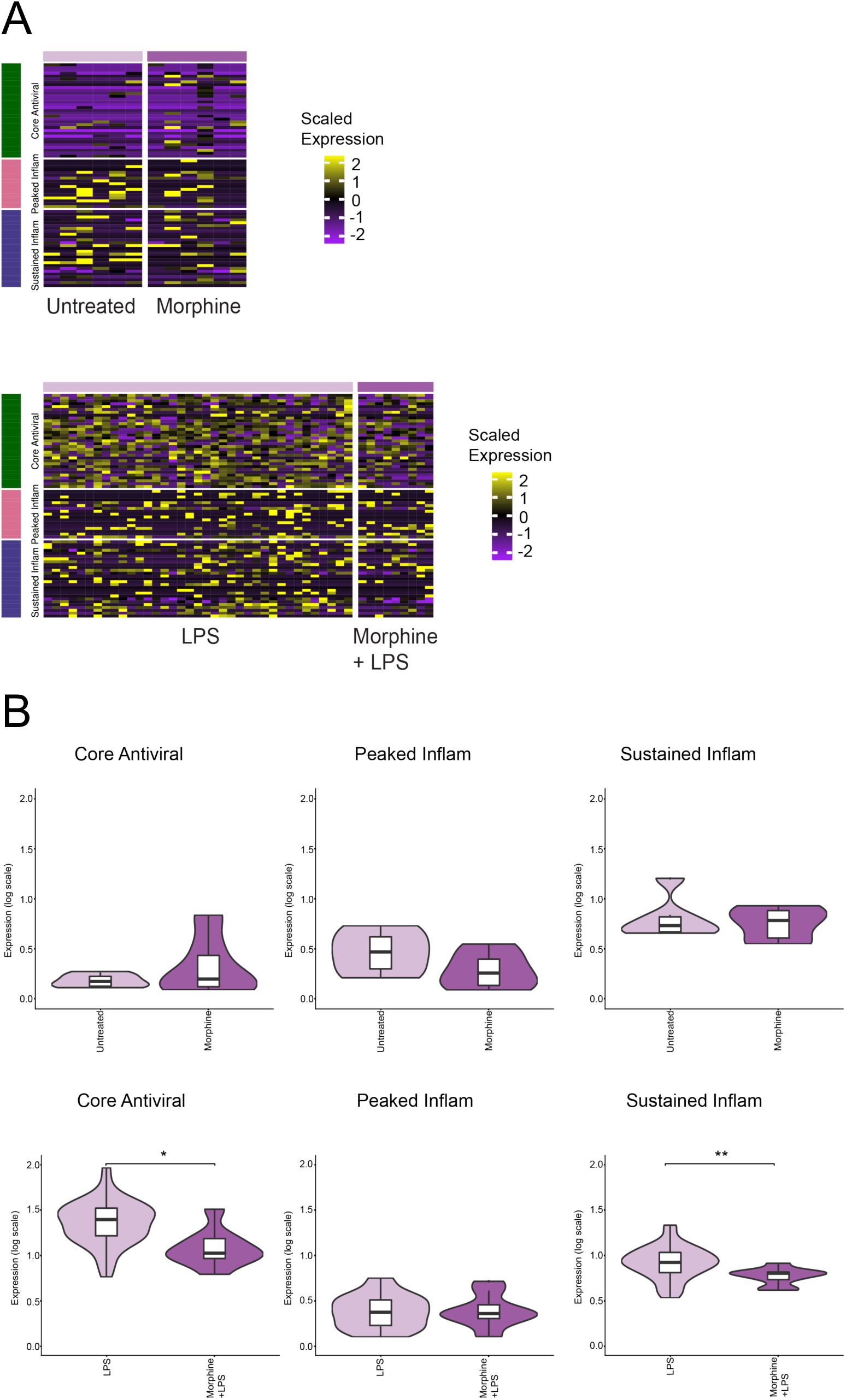
Single cell gene expression heatmap of antiviral and inflammatory gene modules in in vitro treated NK cells. **a,** Heatmap of scaled expression of core antiviral, peaked inflammatory, and sustained inflammatory gene modules observed across all naive state samples (top) and LPS-stimulated samples (bottom). **b,** Average antiviral gene set expression (log expression) across all naive state sample cells (top) and LPS-stimulated sample cells (bottom). Significance shown using p-values: * < 0.05, ** < 0.01, *** < 0.001, **** < 0.0001.

**Figure S17.**
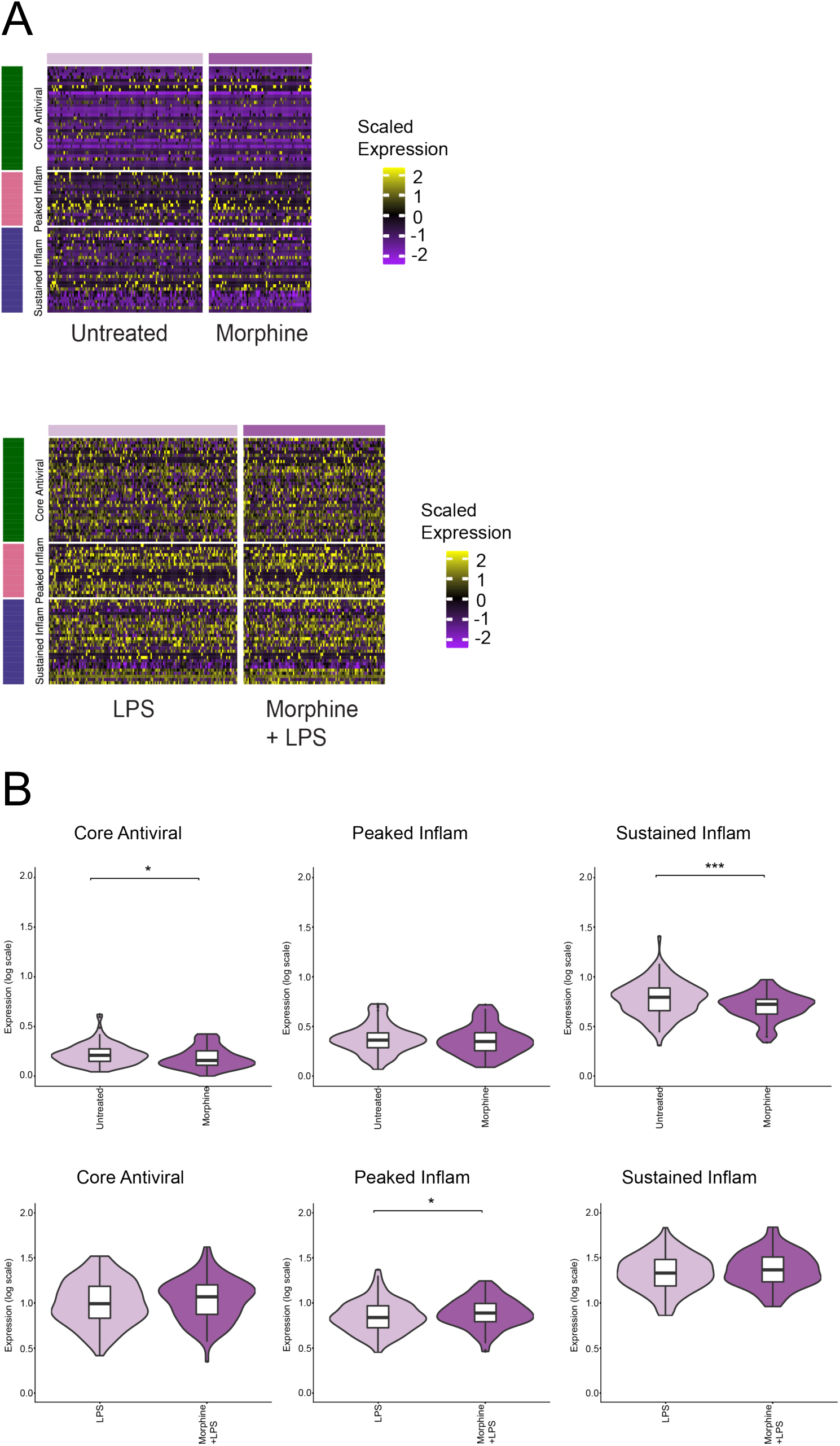
Single cell gene expression heatmap of antiviral and inflammatory gene modules in in vitro treated Monocytes. **a,** Heatmap of scaled expression of core antiviral, peaked inflammatory, and sustained inflammatory gene modules observed across all naive state samples (top) and LPS-stimulated samples (bottom). **b,** Average antiviral gene set expression (log expression) across all naive state sample cells (top) and LPS-stimulated sample cells (bottom). Significance shown using p-values: * < 0.05, ** < 0.01, *** < 0.001, **** < 0.0001.

**Figure S18.**
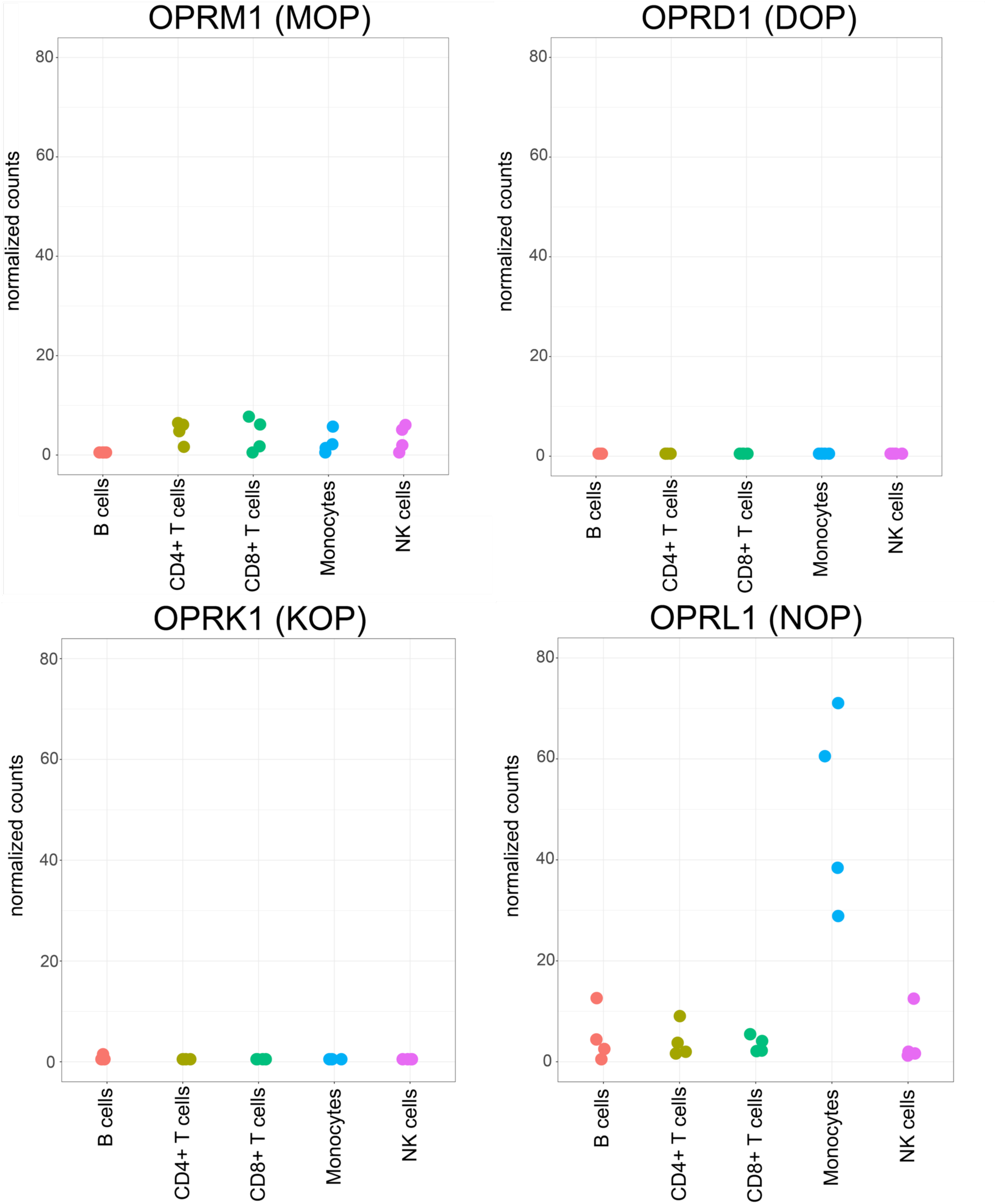
Opioid receptor gene expression in peripheral blood immune cell populations from bulk RNA-seq. Bulk RNA-seq data from Corces et al. 2016^28^ was reanalyzed. Normalized expression of opioid receptor genes MOP (OPRM1), DOP (OPRD1), KOP (OPRK1), and NOP (OPRL1) in healthy control donors across immune cell types: B cells, CD4+ T cells, CD8+ T cells, Monocytes, NK cells.

**Figure S19.**
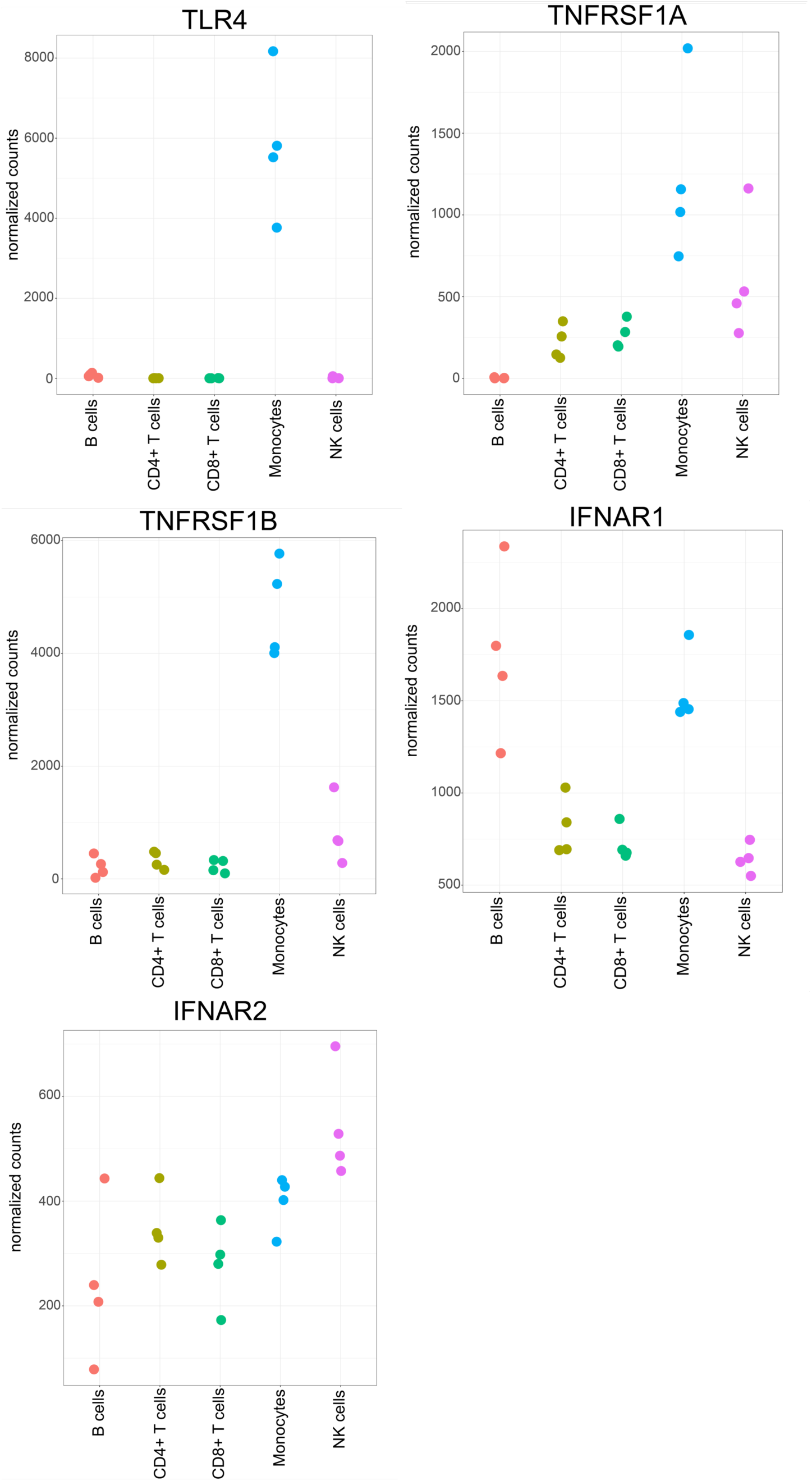
TLR4, TNF receptor and IFN receptor gene expression in peripheral blood immune cell populations from bulk RNA-seq. Bulk RNA-seq data was analyzed from Corces et al. 2016^28^ was reanalyzed. Normalized expression of opioid receptor genes *TLR4*, *TNFRSF1A*, *TNFRSF1B*, *IFNAR1*, *IFNAR2* in healthy control donors across immune cell types: B cells, CD4+ T cells, CD8+ T cells, Monocytes, NK cells.

**Figure S20.**
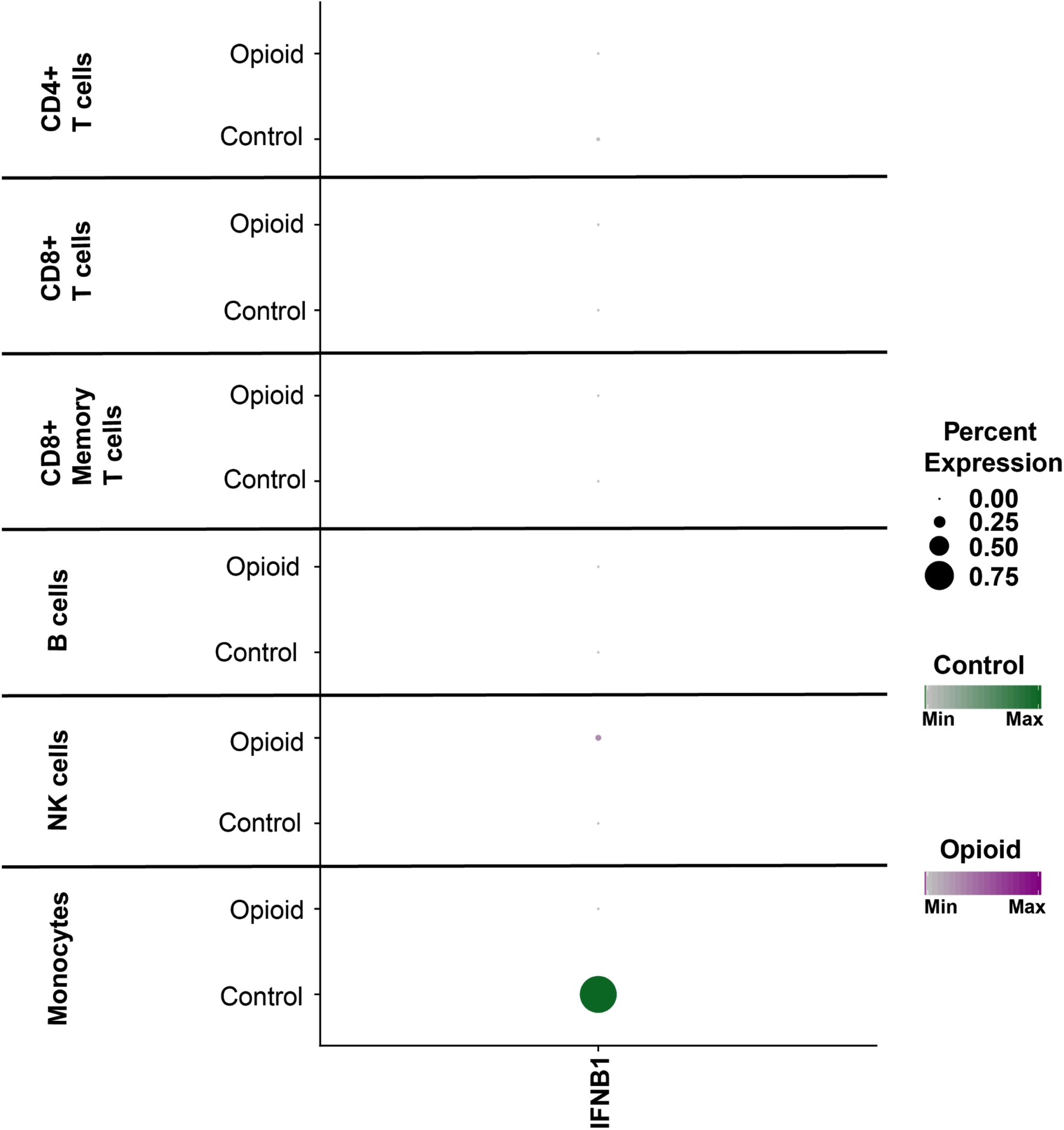
Gene expression of IFNB1 in opioid-dependent and control cells. Dot plot of average gene expression of IFNB1 (x-axis) in control cells (Control) and opioid-dependent cells (Opioid) across naive state cell types: CD4+ T cells, CD8+ T cells, CD8+ Memory T cells, B cells, NK cells, Monocytes (y-axis).

